# Synthetic iminosugar monomers change global metabolic pathways and chitin biosynthesis in *Thalassiosira rotula*

**DOI:** 10.64898/2026.07.09.737548

**Authors:** Jan Ludwig, Timo Watzenborn, Sabine Laschat, Ingrid M. Weiss

## Abstract

- *Thalassiosira rotula* produces extracellular chitin fibers of interest for material science. Tailored monomeric iminosugars, designed as substrate analogues for carbohydrate-active enzymes, unexpectedly elongate these fibers in vivo, yet their impact on chitin metabolism remains unclear.
- *T.rotula* was exposed to three L-isoleucine-derived iminosugar analogues immediately before cell division, when chitin fibers are produced. RNA-sequencing, combined with differential expression and pathway enrichment analyses, as well as transcriptome mining for chitin-related genes was performed.
- Gene mining identified 84 chitin-associated genes (including 42 chitin synthases). Two iminosugars globally repressed carbohydrate- and energy-related pathways including photosynthesis, glycolysis/gluconeogenesis, and Calvin cycle while simultaneously inducing ribosome biogenesis. ImOH specifically downregulated 29 chitin-related genes, including two strongly repressed chitinases and a β-*N*-acetylhexosaminidase.
- Tailored monomeric chitin-modulating iminosugars not only alter chitin fiber length but also trigger a broad metabolic shift from carbohydrate synthesis toward ribosome biogenesis, indicative of a cellular stress response to non-metabolizable iminosugars.

## Introduction

Diatoms (Bacillariophyta) are a class of unicellular, photosynthetic eukaryotes belonging to the diverse group of stramenopiles. They play a major ecological role, accounting for roughly 20% of global annual primary production, sustaining marine food webs and driving the biogeochemical cycling of carbon, nitrogen and silicon (Field *et al*., 1998; Tréguer *et al*., 2018). Their ecological success is tightly linked to the construction of nanopatterned silica cell walls through controlled silicon biomineralization (Martin-Jézéquel, Hildebrand and Brzezinski, 2000; Ehrlich and Witkowski, 2015; Bryłka *et al*., 2024; Safadi *et al*., 2025). Moreover, diatoms are prominent for their distinctive carbon partitioning pathways (Wilhelm *et al*., 2006; Wagner *et al*., 2017). Unlike land plants and green algae, they store carbon mainly as β-1,3-glucans (chrysolaminarin) and triacylglycerols (TAGs) instead of starch (Becker *et al*., 2017). This is a result of a series of endosymbiontic events that provided diatoms with a mosaic genome incorporating genes from green algae, red algae, and cryptophytes, endowing them with these unique metabolic capabilities (Stiller *et al*., 2014; Marella, Bhattacharjya and Tiwari, 2021). Consequently, diatoms possess the alternative glycolytic Entner-Doudoroff pathway and a complete ornithine-urea cycle (OUC) otherwise restricted to metazoans (Bowler *et al*., 2008; Allen *et al*., 2011; Launay *et al*., 2020). The OUC supplies diatoms with a central hub that couples photosynthetic carbon fixation with the synthesis of organic nitrogen compounds, ensuring efficient and robust formation of amino acids and polyamines even under highly fluctuating carbon and nitrogen supply (Allen *et al*., 2011; Prihoda *et al*., 2012). Furthermore, the OUC directly influences glutamine and glutamate synthase cycles which are involved in the formation of key precursors to chitin biosynthesis (Allen *et al*., 2011; Traller *et al*., 2016).

Chitin, the linear polysaccharide of β-1,4 linked *N*-acetyl glucosamine (GlcNAc) units, is the most abundant biopolymer in the marine ecosystem, and its biosynthesis and degradation are pivotal for oceanic biogeochemical carbon and nitrogen cycling (Souza *et al*., 2011; Beier and Bertilsson, 2013). In certain members of the order *Thalassiosirales* (*Thalassiosira* and *Cyclotella*), uniform crystalline chitin fibers are extruded through specialized biosilica pores (fultoportulae) (Blackwell, Parker and Rudall, 1967; Herth, 1979; Lebeau and Robert, 2003; Gügi *et al*., 2015). X-ray diffraction later revealed that these fiber crystals adopt the rare β-allomorph of chitin with a parallel polymer arrangement, in contrast to the antiparallel α-chitin in arthropods (Blackwell, Parker and Rudall, 1967; Wegmann *et al*., 2026). The diatom *Thalassiosira rotula* (synonym of *T.gravida*) is particularly noteworthy, as it produces two distinct chitin fiber populations: larger fibers from central fultoportulae and thinner fibers from peripheral sites, suggesting more elaborate chitin biosynthesis pathways (Ludwig *et al*., 2025). Beyond extracellular fibers which provide diatoms with buoyancy to regulate their position in the water column, chitin is also integrated into the biosilica cell wall, where it contributes to stability and may regulate cell wall integrity (Durkin, Mock and Armbrust, 2009; Wustmann *et al*., 2020).

The advent of high-throughput sequencing has expanded our insight into diatom biology. Genomes and transcriptomes of various diatom species have been released in recent years, allowing metabolic processes to be studied in unprecedented detail (Bowler *et al*., 2008; Dyhrman *et al*., 2012; Shao *et al*., 2019; Cheng, Bowler, *et al*., 2021; Li *et al*., 2021; Downey *et al*., 2023; Di Costanzo *et al*., 2025). Metagenomic and metatranscriptomic surveys of the ocean gene atlas indicate that stramenopiles constitute the second-largest eukaryotic source of chitin synthase transcripts (16% of total hits after crustaceans), with ∼78% of the transcripts derived from *Thalassiosirales*, underscoring the relevance of diatoms in chitin research (Villar *et al*., 2018; Shao *et al*., 2023). Chitin gene mining in *T.weissflogii* identified a complete biosynthetic chitin pathway consisting of a set of 234 different genes (Cheng, Bowler, *et al*., 2021). This extensive repertoire confirms diatoms as attractive models for investigating chitin biosynthesis.

*Thalassiosira rotula* is an attractive model system for quantifying chitin synthase activity in vivo because it synthesizes a linear chitin rod. Roughly 20 µm of chitin rod is synthesized in less than two hours once per cell cycle. Much shorter rods are synthesized in the presence of chitin synthase inhibitors such as Nikkomycin Z in a dose-dependent manner (Schönitzer and Weiss, 2007; Holzwarth *et al*., 2022). In a recent experimental study, supplementation with synthetic L-isoleucine-derived iminosugars modulated chitin fiber synthesis in *T.rotula* in vivo by increasing the length of the produced fibers (Holzwarth *et al*., 2022). Iminosugars are a class of cyclic compounds that feature an endocyclic nitrogen instead of an oxygen atom. These molecules are known as potent inhibitors of carbohydrate active enzymes (CAZy) such as glycosyl hydrolases and glycosyl transferases (Gautier-Lefebvre *et al*., 2005a; Conforti and Marra, 2021) which was contradictory in the case of the *Thalassiosira* study by Holzwarth and colleagues (Holzwarth *et al*., 2022). Although the primary target of iminosugars is likely the active sites of CAZys, transcriptional responses have been documented in bacteria, human macrophages and insects (Rogg *et al*., 2012; Zhang *et al*., 2018; Tomusiak-Plebanek *et al*., 2025). Such responses in diatoms, however, have not yet been reported.

Addressing this gap, the present study leverages the recent functional annotation of *T.rotula* (Di Costanzo *et al*., 2025). Three iminosugars were synthesized, including the compounds that previously elongated chitin fibers. *T.rotula* cells were supplemented with the iminosugars immediately before cell division which is when chitin fiber extrusion starts (Holzwarth *et al*., 2022). RNA-sequencing and differential expression analyses were then performed relative to control cells. Exposure to the two iminosugars ImOH and ImF led to broad repression of carbohydrate- and energy-related pathways including ‘photosynthesis’, ‘glycolysis/gluconeogenesis’, ‘carbon fixation by Calvin cycle’ while ribosome-related processes were upregulated, indicative of a cellular stress response. Moreover, ImOH exposure specifically repressed 29 chitin-related genes, notably two strongly repressed chitinases and a β-*N*-acetyl hexosaminidase.

## Materials and Methods

### Synthesis of a set of iminosugars for transcriptomics

Iminosugars ImOH, and ImCC used in this study were synthesized according to the published protocol (Holzwarth *et al*., 2022). To generate sufficient iminosugar quantities, the synthetic route was scaled up starting from up to 20 g of commercially available L-isoleucine. In a 6-step synthesis, the *N*-protected iminosugars **1** and **2** were synthesized from L-Isoleucine, followed by deprotection with thiophenol to yield the desired iminosugars ImOH and ImCC (Fig. **1**). The fluorinated compound **3** was derived from olefin **1** via a free radical hydrofluorination reaction (Barker and Boger, 2012) with Selectfluor^®^ as F* source and NaBH_4_ as hydride source (Buchholz and Pomerantz, 2021; Gimenez *et al*., 2021; Jones *et al*., 2021; Kumar *et al*., 2025). This method was chosen over the more common functionalization with Olah’s reagent (Py · HF) or other fluoride (F^−^) sources due to the limited scope, as free hydroxyl groups are prone to substitution under these conditions. The hydrofluorination gave fluoride **3** in 53 % yield and successive deprotection under the same conditions with thiophenol led to the desired iminosugar ImF in 70 % yield (Holzwarth *et al*., 2022).

**Figure 1:**
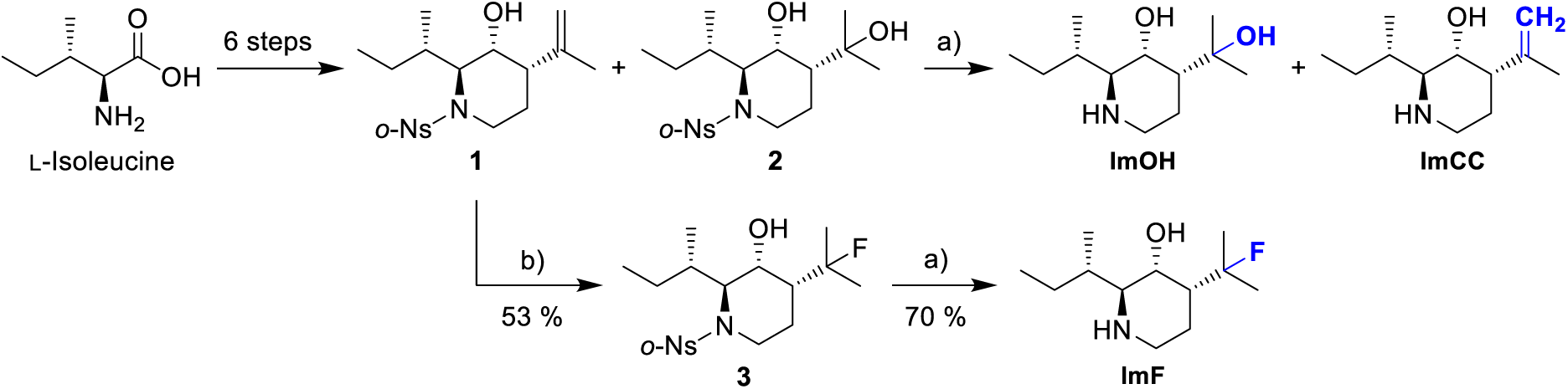
Reagents and conditions: (**a**) PhSH, K_2_CO_3_, DMF, r.t., 24 h, (**b**) Fe_2_(ox)_3_, NaBH_4_, Selectfluor^®^, 0 °C, 30 min. (Barker and Boger, 2012)

### Cultivation of Thalassiosira rotula, experimental design for transcriptomics, and cell harvesting

Cultures of the centric marine diatom *Thalassiosira rotula* were isolated from a marine sample gratefully obtained from the Alfred Wegener Institute in Sylt, Germany. Cultures were maintained in 10 mL culture flasks supplemented with enriched artificial sea water (ESAW) (Harrison, Waters and Taylor, 1980; Berges, Franklin and Harrison, 2001). Before use, the medium was filtered through a 0.2 µm membrane to eliminate contaminants. *T.rotula* was incubated at 18 °C under a light intensity of 50 μmol photons m^−2^ s^−1^ following a 16 h dark: 8 h light cycle (lights on: 6:00 – 14:00).

For transcriptomic experiments, cultures were grown in 10 mL culture flasks until cell density reached 10.000 cells mL^−1^ (∼4-5 days). Cultures were then transferred to 250 mL flasks supplemented with fresh ESAW and grown to the same density (∼4-5 days). A 50 mL aliquot of the running preculture was inoculated into 1950 mL fresh ESAW and allowed to proliferate to 5000 cells mL^−1^. Subsequently, cultures were separated into four 1 L flasks, containing 350 mL cell suspension and 650 mL fresh ESAW. They were maintained until they reached a cell density of 5000 cells mL^−1^ to keep them in exponential growth for the experiments. Just before the time of expected cell division (occurring once per day), each flask was supplemented with iminosugars ImOH, ImF, or ImCC dissolved in 100 µL DMSO to a final concentration of 100 µM in the iminosugar samples or only DMSO for the control. The cells were grown for 5 hours. Each replicate (∼1×10^7^ cells) was harvested by centrifugation at 4000 xg for 4 min at 4 °C. The supernatant was discarded, and the pellet washed three times with phosphate buffered saline (PBS, pH 8.2). After removal of residual buffer, the pellet was flash-frozen in liquid nitrogen and stored at −80°C until RNA extraction.

### Library construction, sequencing and data processing

Total RNA was isolated from *T.rotula* cells and poly-A mRNA was purified at the Novogene facility using poly-T oligo-attached magnetic beads (Novogene LLC, Munich, Germany). RNA integrity was assessed on an Agilent 5400 Bioanalyzer and samples with an RNA integrity score ≥ 4 were retained for library construction. After fragmentation, cDNA synthesis was performed using random hexamer primers, followed by second-strand synthesis, end-repair, A-tailing, adapter ligation, size selection, PCR amplification and purification. Libraries were sequenced in paired-end mode with a read length of 150 nt using the NovaSeq X Plus Series (PE150) (Novogene LLC, Munich, Germany).

Raw reads were trimmed with fastp (v0.23.4) to remove adapter-related reads, poly-*N* stretches, and low-quality bases (Chen *et al*., 2018). Clean reads were mapped against the annotated *T.rotula* transcriptome using HISAT2 (v2.2.1) software with default settings (Kim *et al*., 2019; Di Costanzo *et al*., 2025). Transcript assembly and quantification was carried out with StringTie (v2.2.3) using default parameters (Pertea *et al*., 2015). Gene level read counts were calculated with featureCounts (v2.0.6), and fragments per kilobase of transcript per million mapped reads sequenced (FPKM) were computed for downstream analysis (Liao, Smyth and Shi, 2014).

Differential gene expression analysis employed DESeq2 (v1.42.0) and the *p*-values were adjusted by the Benjamini-Hochberg method (Love, Huber and Anders, 2014). Genes with an adjusted *p*-value (*p*_adj_) ≤ 0.05 and an absolute log₂ fold change |(log_2_FC)| ≥ 1 were deemed significantly differentially expressed. Gene-ontology (GO) enrichment of DEGs was performed with clusterProfiler (v4.8.1) while correcting for gene-length bias (Yu *et al*., 2012). GO terms with *p*_adj_ < 0.05 were considered significant. Kyoto Encyclopedia of Genes and Genomes (KEGG) pathway enrichment was assessed similarly using the same R package. For decluttering GO terms, similar terms were summarized and obsolete terms were discarded using REVIGO (Supek *et al*., 2011).

Gene set enrichment analysis (GSEA) was run with the Broad Institute’s desktop tool (http://www.broadinstitute.org/gsea/index.jsp). Genes were pre-ranked by log_2_FC, and GO and KEGG gene sets were tested independently. Gene sets with a false discovery rate (FDR) < 10% were reported as enriched.

### Hierarchical cluster analysis of chitin related genes

Relative expression of chitin-related genes among different iminosugar treatments were presented as hierarchical clusters using heatmaps. For this, a horizontal hierarchical clustering algorithm was used in NovoMagic (https://eu-magic.novogene.com/) to cluster expression values after log_2_(fpkm+1) transformation and centering correction. Genes were divided into several clusters, and genes within the same cluster show similar expression-level change trends under different treatments.

### Protein sequence analysis and construction of phylogenetic trees

Protein sequences were inferred from the corresponding mRNA transcripts. Subcellular localization predictions employed HEterokont subCellular localization TARgeting (HECTAR; v1.3) (Gschloessl, Guermeur and Cock, 2008). If HECTAR predicted chloroplast localization, ASAFind 2.0 was used to verify the bipartite ASAFAP motif (Gruber *et al*., 2025). Signal peptides were determined with SignalP (v4.1, and v6.0) (Dyrløv Bendtsen *et al*., 2004). Genes were distributed based on HECTAR to type II signal anchors if HECTAR predicted signal anchor, plastids if HECTAR predicted chloroplast and ASAFind 2.0 was positive, mitochondria if HECTAR predicted mitochondrion target peptide, ER if HECTAR predicted signal peptide and cytosol if no signal peptide was detected.

Conserved domains were identified by querying the Pfam (v38.1) database via InterProScan (Jones *et al*., 2014; Paysan-Lafosse *et al*., 2025). Multiple-sequence alignments (MSAs) were performed using Clustal Omega (Sievers *et al*., 2011). Maximum-likelihood neighbor-joining (NJ) trees were constructed in Jalview (v2.11.5.1) using a BLOSUM62 substitution matrix with 1000 bootstrap replicates (Waterhouse *et al*., 2009).

### Quantification and statistical analysis

All experiments were performed in four biological replicates. RNA quantification and statistical analysis are described in the corresponding methodological sections. Statistical analyses were carried out with NovoMagic (https://eu-magic.novogene.com/) and visualizations were generated in R using the ggplot2 package (v4.0.2) (Wickham, 2016).

## Results

### Transcriptome assembly and differential mRNA expression analysis

*T.rotula* cells were grown at constant cell density of 5,000-10,000 cells mL^−1^ and maintained in exponential growth. For each treatment (ImOH, ImF, ImCC) and the DMSO-only control, four RNA-seq libraries were prepared from 10 million cells (40 million cells in total). Cultures received 100 µM iminosugar in 100 µL DMSO before the onset of chitin synthesis, whereas the control received DMSO only. RNA-sequencing generated between 82.6 M and 103.8 M high quality reads (Supporting information Table **S1**). One low quality control library was discarded and excluded from downstream analysis. Mapping to the *T.rotula* reference transcriptome (Di Costanzo *et al*., 2024) (35,230 protein-coding genes, 18909 fully annotated), yielded mapping ratios of 89.5-94.3% (∼77.9 M - 90.7 M total mapped reads per sample) indicating data was sufficient for further analysis. Principal component analysis (PCA) separated the library sets with the four ImOH and the three control replicates forming tight clusters indicating high intragroup similarity (Fig. **S1**). A total of 15222 genes were expressed in all conditions, while 126-193 genes were uniquely expressed (Fig. **S2**) Hierarchical clustering (Fig. **S3**) confirmed intragroup similarity and revealed that ImF and ImOH cluster together, whereas ImCC clusters with the control libraries. In the following, the sets of libraries for each condition are referred to as ImOH, ImF, ImCC, or control.

Differential gene expression (DGE) analysis (|log_2_FC|>1, *p*_adj_<0.05) identified 3957 differentially expressed genes (DEGs) for ImOH, 2219 DEGs for ImF, and 520 DEGs for ImCC relative to the DMSO control. Pairwise comparisons between the iminosugar sets yielded 921 DEGs (ImF vs ImOH), 3721 DEGs (ImCC vs ImOH), and 1490 DEGs (ImCC vs ImF) (Table **1**, Fig. **2a-c**, Fig. **S4**). A Venn diagram (Fig. **2d**) shows that 282 DEGs (compared to control) are shared across all sets. ImOH and ImF share 1794 DEGs, whereas ImOH and ImCC, and ImF and ImCC share only 42 DEGs and 46 DEGs, respectively. Treatment-specific genes comprise 2121 DEGs unique to ImOH, 379 unique to ImF, and 150 unique to ImCC. Detailed information for each DEG is provided in Tables **S2-S5**.

**Figure 2:**
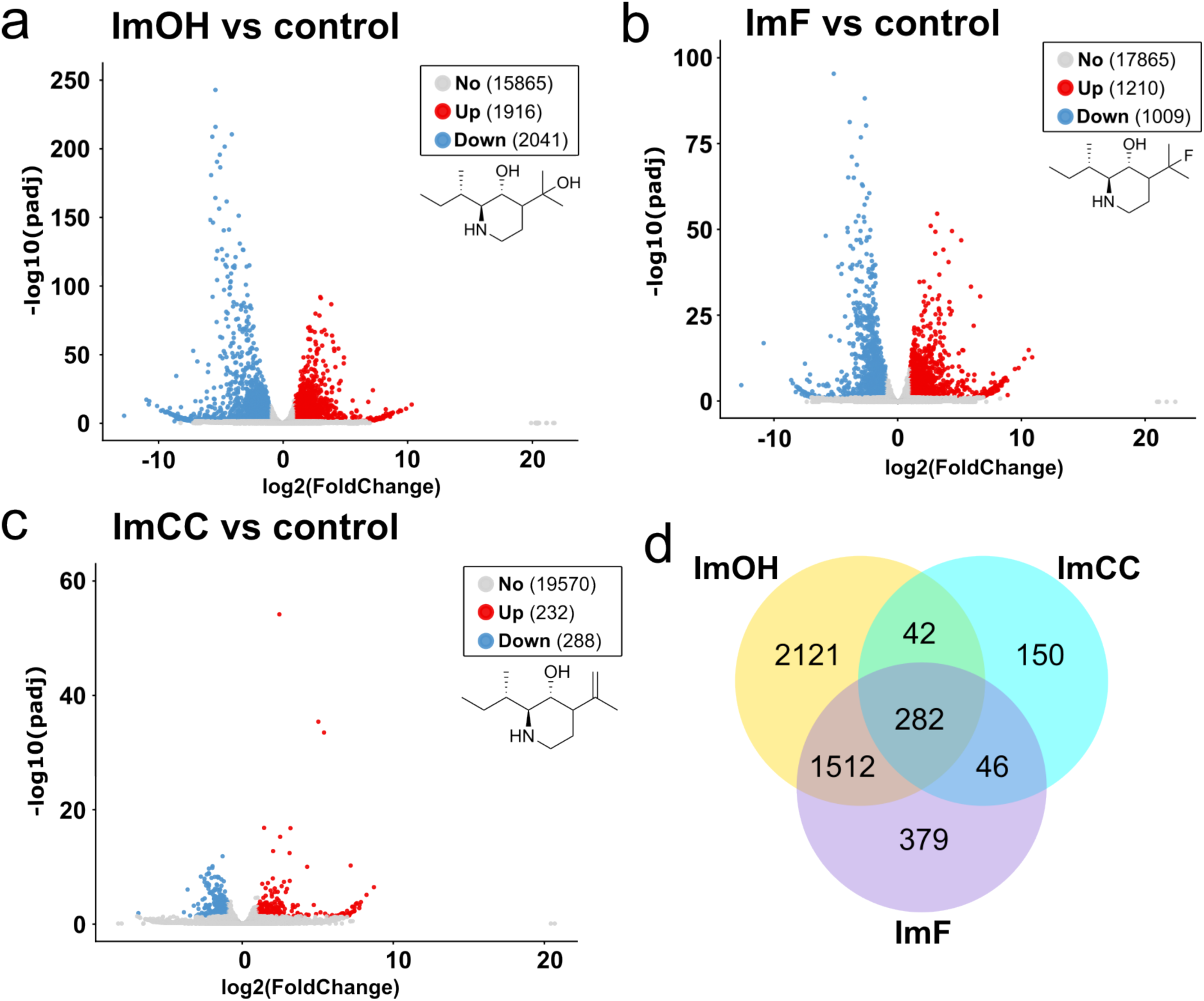
Differentially expressed genes in the iminosugar library sets. Volcano plots show differently expressed genes (|log_2_FC|>1, *p*_adj_ < 0.05) and molecular structure of the iminosugar for (**a**) ImOH vs control, (**b**) ImF vs control, and (**c**) ImCC vs control. Each dot represents a gene, colored by its expression change (gray: not significant, red: upregulated, blue: downregulated). (**d**): Venn diagram shows the overlap between DEGs (relative to the control set) among the three iminosugar library sets. Detailed information for each DEG is found in Tables **S2-S5**.

**Table 1:** Differential gene expression analysis of the iminosugar treatments. (|log_2_FC|>1, *p*_adj_ < 0.05)

| Comparison | Total DEGs | Up-regulated | Down-regulated | Non DEG |
| --- | --- | --- | --- | --- |
| ImOH vs control | 3957 | 1916 | 2041 | 15865 |
| ImCC vs control | 520 | 232 | 288 | 19570 |
| ImF vs control | 2219 | 1210 | 1009 | 17865 |
| ImCC vs ImOH | 3721 | 2124 | 1597 | 16401 |
| ImF vs ImOH | 921 | 733 | 188 | 19098 |
| ImCC vs ImF | 1490 | 654 | 836 | 18789 |

### Functional enrichment analysis of differentially expressed genes

KEGG pathway enrichments of the DEGs (|log_2_FC|>1, *p_adj_* < 0.05) revealed distinct transcriptional profiles for the three iminosugars (Fig. **3**). In ImOH, 21 pathways were significantly altered (Fig. **3a**). Although 1916 of the 3957 total DEGs were upregulated, only three pathways were enriched among them (‘ribosome’, ‘ribosome biogenesis in eukaryotes’, and ‘phenylalanine, tyrosine, and tryptophan biosynthesis’. Corresponding gene ontology (GO) terms in the biological processes (BP) aspect (‘peptide biosynthetic process’, ‘peptide metabolic process’, and ‘translation’) and molecular function (MF) aspect (‘structural constituent of ribosome’, ‘catalytic activity on RNA’, ‘catalytic activity on tRNA’ and ‘aminoacyl tRNA ligase activity’) ranked among the most significant (Fig. **S5**, Table **S6**). ImF showed a comparable pattern, with 17 significantly enriched pathways in total (Fig. **3b**).

**Figure 3:**
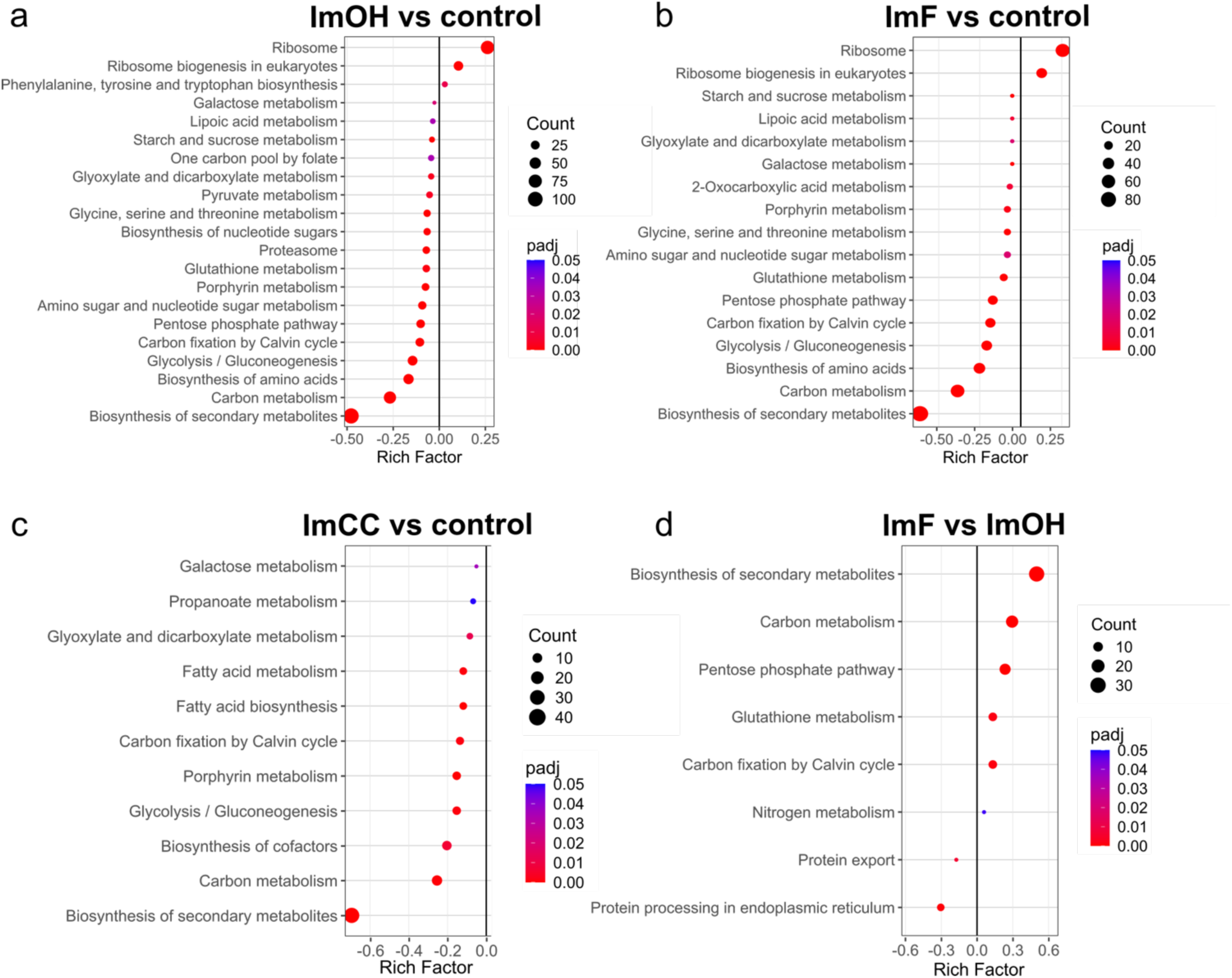
KEGG enrichment analysis of differently expressed genes (DEG)s (|log_2_FC|>1, *p*_adj_ < 0.05) under different iminosugar treatments. (**a**) ImOH vs control, (**b**) ImF vs control, (**c**) ImCC vs control. (**d**) ImF vs ImOH. The rich factor is depicted on the x-axis. Pathways enriched among downregulated DEGs are signified by negative and upregulated DEGs by positive rich factors. A list of the genes of each KEGG pathway enrichment is listed in Table **S7**.

Upregulated DEGs were enriched for ‘ribosome’, and ‘ribosome biogenesis in eukaryotes’ pathways. By contrast, ImCC displayed no significant enrichment among upregulated DEGs and showed only 11 downregulated pathways (Fig. **3c**).

Across all three iminosugar sets, downregulated DEGSs were consistently enriched in carbohydrate-and energy-metabolism pathways such as ‘glycolysis/gluconeogenesis’, ‘carbon fixation by the Calvin cycle’, and general ‘carbon metabolism’ (Fig. **3a-c**). ImOH and ImF additionally showed repression of the ‘pentose phosphate pathway’, ‘aminosugar and nucleotide sugar metabolism’, ‘starch and sucrose metabolism’, ‘glutathione metabolism’ and the ‘porphyrin metabolism’ (photosynthesis) (Table **S7**). GO-BP terms related to photosynthesis (‘photosynthesis’, ‘tetrapyrrole biosynthetic process’, ‘tetrapyrrole metabolic process’) ranked among the top enriched for ImCC and appeared at lower significance for ImOH and ImF (Table **S6**). Uniquely for ImCC, downregulated DEGs were enriched in ‘fatty acid metabolism’ and ‘fatty acid biosynthesis’ pathways (Fig. **3c**).

Relative to the DMSO-control, both iminosugar sets showed significant enrichment of upregulated genes in the ribosomal KEGG pathways ‘ribosome’ and ‘ribosome biogenesis in eukaryotes’. Because of this, ImOH was contrasted against ImF (Fig. **3d**). No significant difference in these enrichments is observed. However, ImF exhibits significantly stronger enrichment of ‘carbon metabolism’, ‘pentose phosphate pathway’, and ‘carbon fixation by Calvin cycle’ among upregulated genes, indicating weaker overall repression of the carbohydrate metabolism compared to ImOH. Conversely, downregulated genes are more enriched for ‘protein export’ and ‘protein processing in endoplasmatic reticulum’ in ImF.

Pre-ranked gene set enrichment analysis (GSEA) confirmed DGE results, including enrichment of ‘ribosome biogenesis in eukaryotes’ and ‘Phe- Tyr- and Trp-biosynthesis’ in ImOH, whereas key carbohydrate pathways, such as ‘carbon metabolism’, and ‘glycolysis/gluconeogenesis’ were enriched in the control (Table **S8**).

### Upregulation of ribosomal processes under iminosugar treatment

Next, ribosome-related transcription was analyzed (Fig. **4**). ImOH and ImF displayed 126 and 97 ribosome-related DEGs, of which 122 and 96, were significantly upregulated (Table **S9**, **S10**). This upregulation encompassed genes encoding for tRNA ligases, 90S pre-ribosomal factors (UTP-B, UTP-C, tUTP, MPP10), rRNA modifying enzymes (rRNA 2’-O-methyltransferases, pseudouridine synthases), nucleolar export proteins, and pre-40S /pre-60S maturation factors (Fig. **4a**). Moreover, expression of structural ribosomal proteins was upregulated (ImOH: 43 large subunit and 30 small subunit proteins, ImF: 29 large and 23 small subunit proteins, Fig. **4b**). Overall, these findings show broad activation of the ribosome machinery and a shift towards protein synthesis in ImOH and ImF.

**Figure 4:**
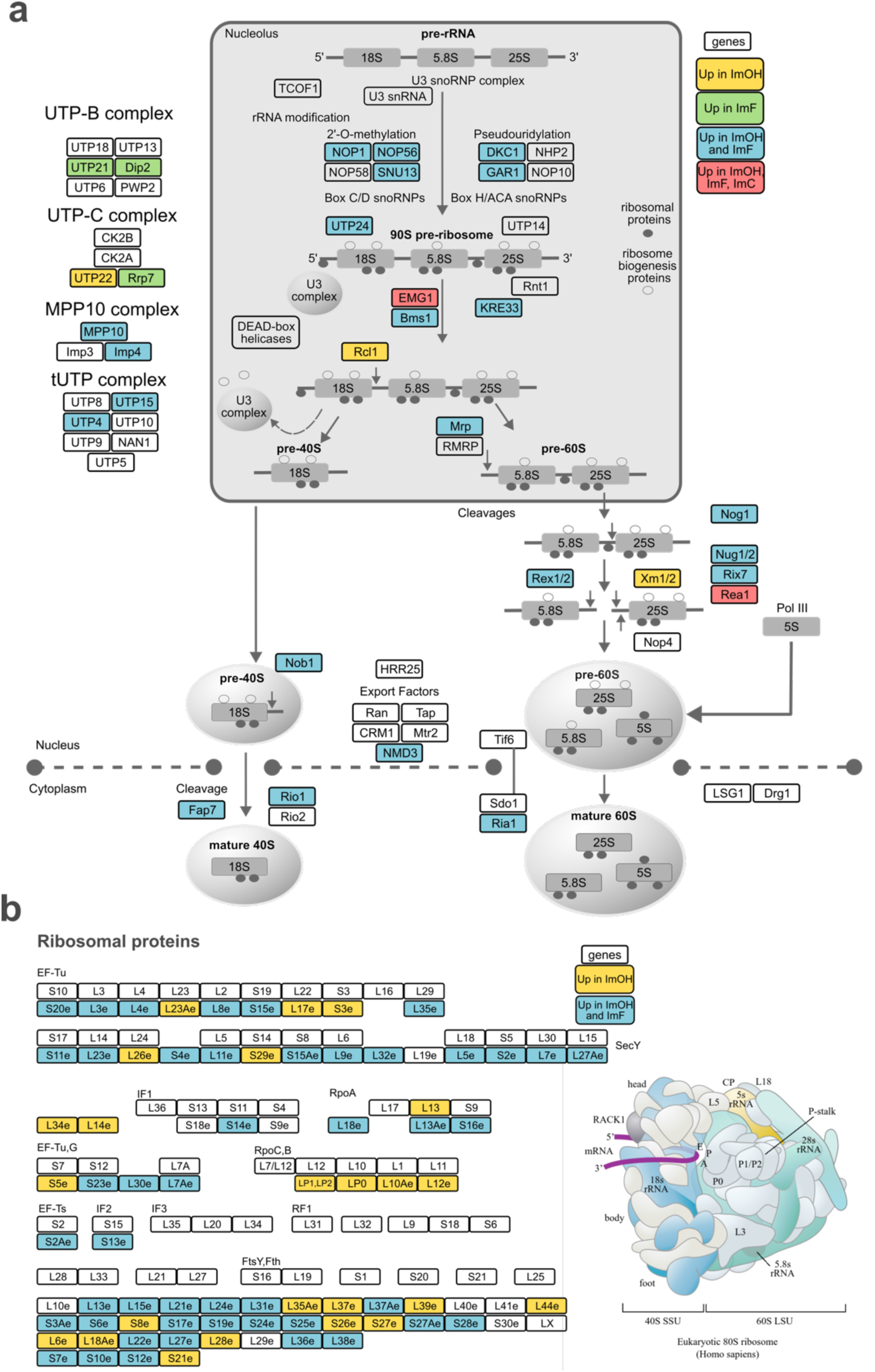
Upregulated genes of ribosome related pathways in ImOH, ImF, and ImCC library sets. (**a**) KEGG map03008 (ribosome biogenesis in eukaryotes) and (**b**) KEGG map03010 (ribosome). Genes with log_2_FC>1, *p*_adj_ < 0.05 are highlighted. Yellow = exclusively upregulated in ImOH, green = exclusively upregulated in ImF, blue = upregulated in both ImOH and ImF, and red = upregulated in ImOH, ImF, and ImCC.

### Global shift of key metabolic pathways under iminosugar treatment

To identify transcriptional changes affecting the chitin metabolism, we examined pathways producing UDP-*N*-acetylglucosamine (UDP-GlcNAc)-precursor fructose-6-phosphate (F6P), namely ‘photosynthesis’, ‘carbon fixation by Calvin cycle’, ‘glycolysis/gluconeogenesis’, and the TCA cycle. Additionally, because a glutamine donor is required for UDP-GlcNAc synthesis, the nitrogen metabolism and ornithine-urea-cycle (OUC) which is connected to glutamine/glutamate cycle, were examined. Fig. **5** displays the DEGs in ImOH (vs control) and highlights the key pathway interconnections. (DEG counts per pathway are provided in Table **2**; full gene list in Supplementary Table **S9-S11**)

**Figure 5:**
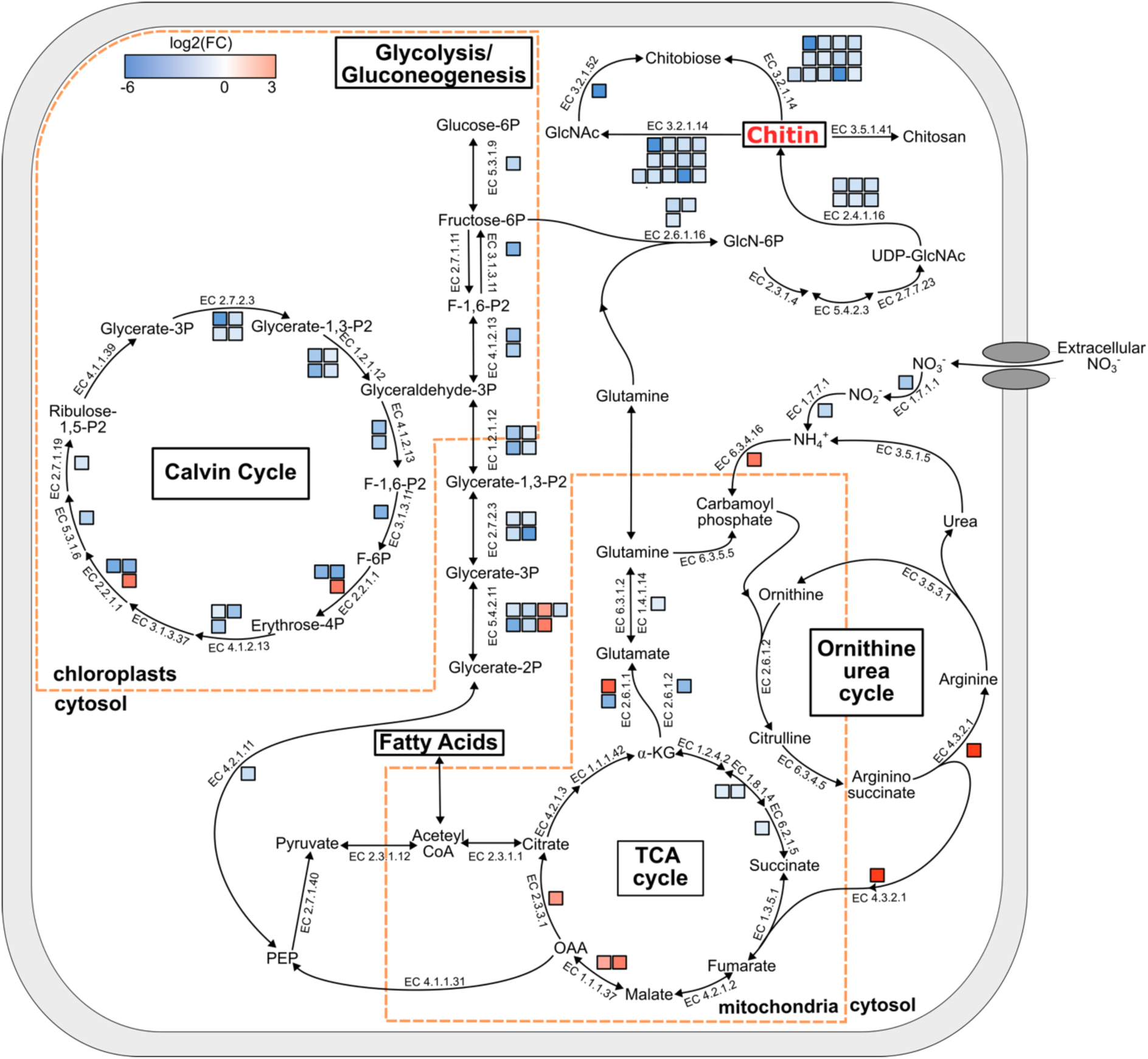
Differentially expressed genes (DEG) in ImOH (vs control) across key metabolic pathways for chitin metabolism. Each box represents a single DEG (|log_2_FC|>1, *p*_adj_ < 0.05) colored according to its log_2_FC. Dashed outlines indicate different subcellular compartments while rectangles show corresponding metabolic pathways. The schematic layout follows the scaffold presented by (Downey *et al*., 2023).

**Table 2:** Number of differentially expressed genes for each pathway contributing to chitin metabolism.

| Pathway | ImOH |  | ImF |  | ImCC |  |
| --- | --- | --- | --- | --- | --- | --- |
|  | Up | Down | Up | Down | Up | Down |
| Photosynthesis | 10 | 111 | 10 | 100 | 1 | 17 |
| Glycolysis / Gluconeogenesis and carbon fixation by Calvin cycle | 7 | 45 | 8 | 41 | 1 | 13 |
| TCA cycle | 3 | 6 | 5 | 3 | 0 | 1 |
| Nitrogen metabolism / Urea-Ornithine cycle | 3 | 6 | 3 | 5 | 1 | 2 |

#### Photosynthesis

Expression of photosynthetic genes was predominantly repressed, with 111 and 100 downregulated DEGs in ImOH and ImF, respectively, versus only 17 in ImCC (Table **2**, Table **S9-S11**). The majority of the downregulated transcripts belong to diatom-specific fucoxanthin chlorophyll *a*-*c* binding proteins (FCPs) with 80 DEGs (mean log_2_FC = −4.71) for ImOH, 73 (mean log_2_FC = −2.45) for ImF and 4 (mean log_2_FC = −2.24) for ImCC. One FCP in ImOH was upregulated (log_2_FC = 3.47). Additionally, strongly downregulated DEGs included genes encoding chlorophyll a-b binding protein LI818 (log_2_FC = −6.85 in ImOH, −5.15 in ImF), ferredoxin 1 (log_2_FC = −4.54 in ImOH, −4.55 in ImF), and components of photosystem II such as Psb31 (log_2_FC = −4.45 in ImOH, and −2.68 in ImF), HCF136 (log_2_FC = −4.41 in ImOH, −4.25 in ImF, −2.23 in ImCC) and Pbs27 (log_2_FC = −3.68 in ImOH, −3.02 in ImF, −1.53 in ImCC). The most upregulated genes in ImOH encoded a ferredoxin-1 (log_2_FC = 3.14) and a chlorophyll a/b-binding protein 5 (log_2_FC =2.02). Similar upregulation of a ferredoxin 1-coding gene was observed in ImF (mean log_2_FC =2.44) and ImCC (log_2_FC =1.90).

#### Calvin cycle, glycolysis/gluconeogenesis, and TCA cycle

F6P is produced from glyceraldehyde-3-phosphate (G3P) either as a Calvin cycle intermediate or via gluconeogenesis from phosphoenolpyruvate (PEP) (Fig. **5**). Across the Calvin cycle and glycolysis/gluconeogenesis, broad transcriptional repression was observed. In total, 52 DEGs in ImOH, 49 in ImF, and 14 in ImCC were differently expressed of which 45, 49, and 14, were downregulated, respectively (Table **2**). Repressed Calvin cycle DEGs included genes encoding for phosphoglycerate kinases (EC 2.7.2.3, mean log_2_FC = −2.43 in ImOH, −1.70 in ImF, −1.18 in ImCC), glyceraldehyde-3-phosphate dehydrogenases (EC 1.2.1.12, mean log_2_FC = −2.50 in ImOH, −2.00 in ImF, log_2_FC = −1.45 in ImCC), and fructose-bisphosphate aldolases (EC 4.1.2.13, mean log_2_FC = −2.97 in ImOH, and −2.02 in ImF). In ‘glycolysis/gluconeogenesis’, transcription of enolase (EC 4.2.1.11, log_2_FC = −1.83 in ImOH, −1.26 in ImCC) was repressed. Seven DEGs encoding 2,3-bisphosphoglycerate-dependent phosphoglycerate-mutase (EC 5.4.2.11) were identified in ImOH, of which five were down- (mean log_2_FC = −2.33) and two upregulated (mean log_2_FC = 1.55). ImF contained four down- (mean log_2_FC = −2.17) and one upregulated DEG (log_2_FC = 1.52), while ImCC showed a single downregulated DEG (log_2_FC = −2.02). No DEGs were detected for chrysolaminarin synthesis, the main carbohydrate storage glucan in diatoms, but across all DEGs, six glucanases were found down- (mean log_2_FC = −2.19) and one upregulated (log_2_FC = 2.74) in ImOH.

Analysis of TCA cycle genes revealed nine DEGs in ImOH, eight in ImF, and one in ImCC. Genes encoding malate dehydrogenase (EC 1.1.1.37, mean log_2_FC = 1.50 in ImOH, 1.71 in ImF) and citrate synthase (EC 2.3.3.1, log_2_FC = 1.42 in ImOH, and 2.00 in ImF) were consistently upregulated, suggesting increased flux towards citrate and downstream fatty acid synthesis. In ImOH, two genes encoding acetyl-CoA acyltransferases (EC 2.3.1.16, mean log_2_FC = 1.15) and a long chain fatty acid CoA ligase (EC 6.2.1.3, log_2_FC = 2.19) were upregulated. Moreover, an upregulated DEG encoding elongation of very long chain fatty acids protein 4 was identified (EC 2.3.1.199, log_2_FC =1.29 in ImOH, log_2_FC =1.77 in ImF). Furthermore, a gene coding for diacylglycerol acyltransferase (DGAT) (EC 2.3.1.20, log_2_FC = 3.46) was found upregulated in ImOH. DGAT is known to catalyze the conversion of diacylglycerol to triacylglycerols (TAGs) which are the major storage lipid in diatoms (Murison *et al*., 2025).

#### Nitrogen metabolism and ornithine urea cycle (OUC)

UDP-GlcNAc synthesis requires the transfer of glutamine onto F6P. In cells, glutamine and glutamate can be interconverted by glutamine synthases (EC 6.3.1.2, and 1.4.1.14) and glutaminases (EC 3.5.1.2) linking nitrogen assimilation and transport to both the TCA cycle and OUC (Fig. **5**). Across the iminosugar libraries, DEGs involved in nitrogen metabolism and the OUC were detected (9 in ImOH, 8 in ImF, 3 in ImCC; Table **2**). Genes encoding nitrate reductase (EC 1.7.1.1, log_2_FC =-2.87 in ImOH, −1.27 in ImCC) and nitrite-reductase (EC 1.7.7.1, log_2_FC = −2.10 in ImOH, −2.29 in ImF, −1.25 in ImCC) were downregulated, whereas, a gene encoding carbamoyl phosphate synthase (EC 6.3.4.16) was found upregulated in ImOH (log_2_FC = 1.86). A gene coding for argininosuccinate lyase (EC 4.3.2.1) showed similar induction (log_2_FC = 2.68 in ImOH, 2.50 in ImF), indicating a shift from extracellular nitrogen uptake towards incorporation of intracellular ammonia into carbamoyl phosphate and subsequent amino acid synthesis.

### Chitin gene mining and chitin metabolism

From KEGG map00520 ‘amino-sugar and nucleotide sugar metabolism’ and targeted query of the annotated transcriptome for ‘UDP-*N*-acetylglucosamine transporter’ and ‘chitin binding domains’ (Table **3**), 84 expressed genes associated with chitin metabolism were retrieved and grouped into three functional classes. Chitin anabolism containing 48 genes including six glutamine-fructose-6-phosphate amidotransferase (GFAT), and 42 chitin synthases (CHS), chitin catabolism containing 26 genes including 23 chitinases, and three β-*N*-acetylhexosaminidase (NAHase), and other chitin-related proteins which contain 10 genes including three chitin binding domain (CBD)-containing proteins, and seven UDP-GlcNAc transporters. Absence of transcripts for other enzymes involved in chitin biosynthesis likely reflects incomplete annotation. Differential expression analysis revealed DEGs in each class combined across all libraries (three GFATs, eight chitin synthases, one NAHase, 13 chitinases, two CBDs, three UDP-GlcNAc transporters) with the majority found in ImOH (three GFATs, six CHS, one NAHase, 13 chitinases, two CBDs, three GlcNAc transporters). Five DEGs were found in ImF (two CHS, three chitinases), and no significant DEG in ImCC.

**Table 3:** Enzymes related to chitin.

|  | EC number | Annotated entries | DEGs across libraries |
| --- | --- | --- | --- |
| **Anabolism** |  |  |  |
| Glutamine-fructose-6-phosphate amidotransferase (GFAT) | 2.6.1.16 | 6 | 3 |
| Glucosamine-phosphate *N*-acetyltransferase | 2.3.1.4 | 0 | 0 |
| Phosphoacetylglucosamine mutase | 5.4.2.3 | 0 | 0 |
| UDP-*N*-acetylglucosamine diphosphorylase | 2.7.7.23 | 0 | 0 |
| *N*-acetylglucosamine kinase | 2.7.1.59 | 0 | 0 |
| Chitin synthase (CHS) | 2.4.1.16 | 42 | 8 |
| **Catabolism** |  |  |  |
| Chitinase | 3.2.1.14 | 23 | 13 |
| β-*N*-acetylhexosaminidase (NAHase) | 3.2.1.52 | 3 | 1 |
| Chitin Deacetylase | 3.5.1.41 | 0 | 0 |
| **Other** |  |  |  |
| Chitin binding domain containing proteins | - | 3 | 2 |
| *N*-acetylglucosamine transporter | - | 7 | 3 |

A hierarchical cluster heat map of row-wise z-scores was constructed and shows the relative expression levels among each chitin-related gene across treatments (Fig. **6**). ImOH and ImF sets cluster closely together while ImCC clusters with the control, mirroring the whole transcriptome profile (Fig. **S3**). All chitin-related genes were downregulated except TROT027331 which codes for an UDP-GlcNAc transporter in ImOH (log_2_FC = 1.23). Among the seven GFAT transcripts, three (TROT016862, TROT011196, TROT011836) were significantly repressed in ImOH (mean log_2_FC = −1.58). Of 42 genes encoding CHS, six were downregulated in ImOH (mean log_2_FC = −1.45) and two (TROT025394, TROT024243) exclusively in ImF (mean log_2_FC = −1.36). One NAHase gene (TROT028691) was strongly downregulated in ImOH (log_2_FC = −7.36). 13 of 25 genes encoding chitinases were downregulated in ImOH (mean log_2_FC = −2.39), including strongly repressed TROT011391 (log_2_FC = −6.92) and TROT031324 (log_2_FC = −7.11). Three of these (TROT008789, TROT005249, TROT011391) were also downregulated in ImF (mean log_2_FC = −3.00). Two genes encoding for CBD-containing proteins (TROT000856, TROT003885, mean log_2_FC = −2.73) and two additional UDP-GlcNAc transporters (TROT010223, TROT033515, mean log_2_FC = −1.59) were downregulated in ImOH.

**Figure 6:**
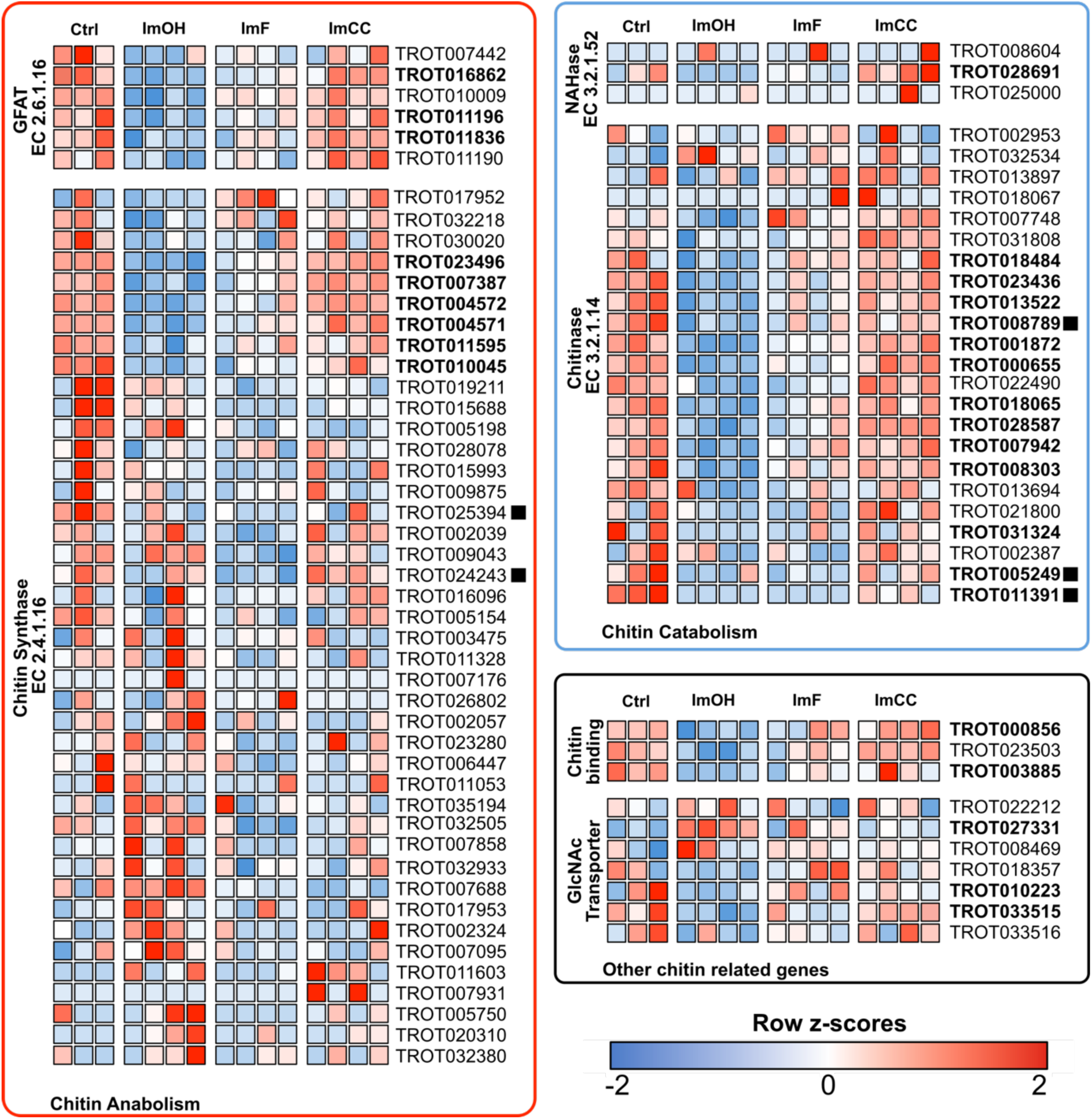
Hierarchical cluster analysis of chitin-related genes in *T.rotula* across the libraries. Each row represents one gene (ID shown on the right). Each box represents the row z score for a single replicate. Genes that satisfy the criteria for differential expression (|log_2_FC|>1, *p*_adj_ < 0.05) are emphasized by bold gene IDs for ImOH or black squares next to the gene ID for ImF.

To investigate the structural features and subcellular distribution of the ImOH DEGs, HECTAR was employed for localization prediction (Fig. **7a**) and InterProScan for domain analysis (Fig. **7b-g**, **S6**).

**Figure 7:**
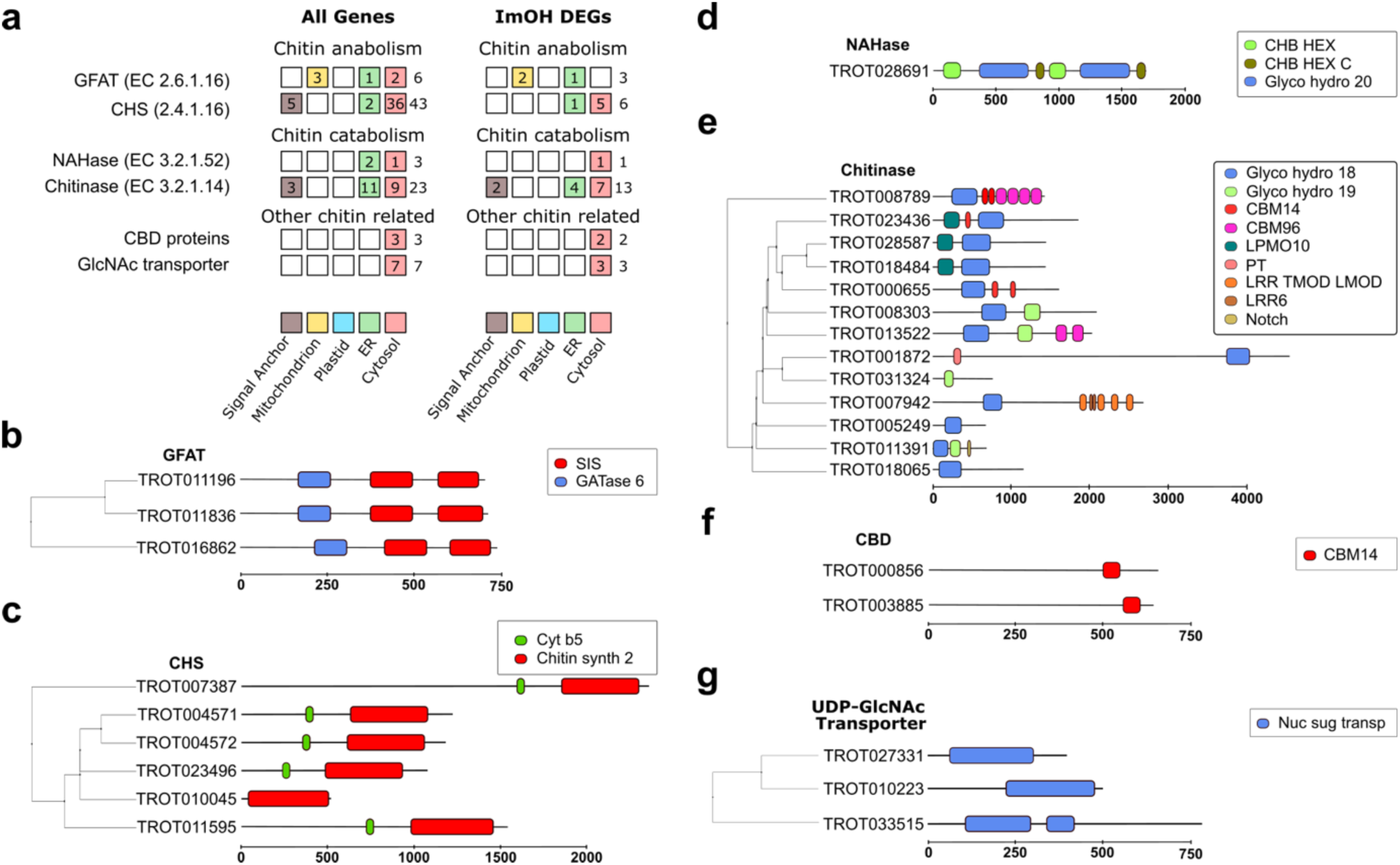
Predicted subcellular localization and domain architectures of differentially expressed chitin-related genes for ImOH vs control. (**a**) Predicted subcellular localization of chitin related genes (**b**) glutamine fructose amidotransferase (GFAT), (**c**) chitin synthase (CHS), (**d**) N-acetyl hexosaminidase (NAHase), (**e**) chitinase, (**f**) chitin binding domain (CBD)-containing proteins and (**g**) UDP-GlcNAc transporter.

Among all GFATs in the *T.rotula* transcriptome, three were predicted to localize to the mitochondria, one to the endoplasmatic reticulum (ER) and two to the cytosol. GFAT DEGs included two mitochondrial and one cytosol-targeted, each bearing a glutamine amidotransferase (GATase 6, pfam13522) domain together with two upstream sugar isomerase (SIS, pfam01380) domains (Fig. **7b**). Five CHS were predicted to contain a type II signal anchor (SA type-II), two were ER-targeted and 35 cytosolic. Of the DEGs, five were cytosolic, all of which contained a cytochrome b5 domain (cyt b5, pfam00173) and the one lacking the cyt-b5 domain was ER-targeted. Strikingly, cyt b5 domains seem remarkably common among the CHS sequences as they were present in 35 of 42 across all libraries (Fig. **S6**). Moreover, each CHS contained the chitin synthase family 2 domain (chitin synth 2, pfam03142, Fig. **7c**). NAHases included two ER-targeted and one cytosolic protein, the latter being the DEG TROT028691 (log_2_FC= −7.36) exhibiting a dual repeat of CHBHex (pfam03173), a glycosyl hydrolase family 20 (pfam00728) and a CHBHexC (pfam03174) domain (Fig. **7d**). The chitinase set consisted of three SA type-II, 11 ER-localized, and nine cytosolic proteins. Of the 13 chitinase DEGs, two have SA type-IIs (TROT001872, TROT023436), four are ER-localized (TROT000655, TROT007942, TROT018065, TROT018484) and seven are cytosolic including the strongly downregulated TROT011391 (log_2_FC= −6.92) and TROT031324 (log_2_FC= −7.11). The domain architecture of the chitinase DEGs (Fig. **7e**) was diverse displaying nine conserved domains including the common glycosyl hydrolase family 18 (GH18, pfam00704) and the rarer family 19 (GH19, pfam00182) domain. The GH18 domain appeared exclusively in nine DEGs, while GH19 appeared in TROT031324. Three DEGs (TROT00803, TROT013522, TROT011391) exhibited a dual architecture comprising both family 18 and 19 domains. Interestingly, the two strongly downregulated chitinases TROT011391 (log_2_FC= −6.92) and TROT031324 (log_2_FC= −7.11) both include GH19 domains. The chitinase domain repertoire was further diversified by lytic polysaccharide monooxygenase (LPMO 10, pfam03067) domains in three DEGs (TROT023436, TROT028587, and TROT018484), chitin binding peritrophin-A domains (CBM 14, pfam01607) in three sequences (TROT008789, TROT023436, 000655), carbohydrate binding module family 6 (CBM 96, pfam24517) in two sequences (TROT008789, TROT013522), PT disordered repeats (pfam04886) in TROT001872, and leucine rich repeats (pfam13516) combined with tropomodulin and leiomodin (TMOD/LMOD, pfam27087) in TROT007942. An LNR domain (Notch, pfam00066) was identified in TROT011391. All CBD-containing proteins and UDP-GlcNAc transporters were predicted to localize to the cytosol (Fig. **7a**). The two CBD-containing protein DEGs (Fig. **7f**) each contained a CBM 14 domain, whereas every UDP-GlcNAc transporter (Fig. **7g**) contained a nuclear sugar transporter domain (Nuc sug transp, pfam04142).

## Discussion

Chitin, the β-1,4-linked GlcNAc polymer, is the most abundant marine polysaccharide and has a key role in oceanic biogeochemical carbon- and nitrogen cycling (Souza *et al*., 2011; Beier and Bertilsson, 2013). In diatoms, chitin reinforces the silica cell wall termed the frustule (Durkin, Mock and Armbrust, 2009; Wustmann *et al*., 2020) and in several centric species, it is extruded as pure β-chitin fibers through specialized fultoportulae (Blackwell, Parker and Rudall, 1967; Herth, 1979; Lebeau and Robert, 2003; Gügi *et al*., 2015). In some diatoms, chitin can make up 31-38% of total dry cell mass indicating it might be a significant carbon reservoir in the cell next to chrysolaminarin and triacylglycerols (McLachlan, McInnes and Falk, 1965).

### Chitin related gene mining in Thalassiosira rotula

Using the recently released genome annotation (Di Costanzo *et al*., 2025), detailed gene mining in *T.rotula* identified 84 expressed chitin-related genes (Table **3**), of which 48 are involved in chitin biosynthesis and 26 in degradation. Compared to other diatoms (*Thalassiosira weissflogii* (234), *Cyclotella nana* (synonym of *Thalassiosira pseudonana*) (141), *Phaeodactylum tricornutum* (48) and *Cyclotella cryptica* (50); Table **4**) *T.rotula* has an intermediate chitin gene count (Traller *et al*., 2016; Cheng, Bowler, *et al*., 2021). The apparent lack of *N*-acetyl glucosamine-6-phosphate deacetylase, phosphoacetyl glucosamine mutase, and UDP-*N*-acetyl glucosamine diphosphorylase transcripts likely reflects incomplete annotation as these genes are scarce also in other diatoms. A striking feature of *T.rotula* is the high number of CHS transcripts (42) exceeding those reported of *T.weissflogii* (30), *C.nana* (20), *P.tricornutum* (2), or *C.cryptica* (7). This might indicate specialization of different CHS isoforms for distinct fultoportulae, consistent with the observation that chitin fiber morphology varies with extrusion site (Ludwig *et al*., 2025). Only 23 different chitinase transcripts were found in *T.rotula*, which is comparable to *C.cryptica* (22), but fewer than in *T.weissflogii* (124), *C.nana* (61), or *P.tricornutum* (36). The number of NAHase transcripts in *T.rotula* is similar to that of *C.nana* (1), *P.tricornutum* (2), and *C.cryptica* (2) but lower than that of *T.weissflogii* (17). Overall, the low number of catabolic (26) compared to anabolic transcripts (48) in *T.rotula* suggests bias towards chitin biosynthesis.

**Table 4:**
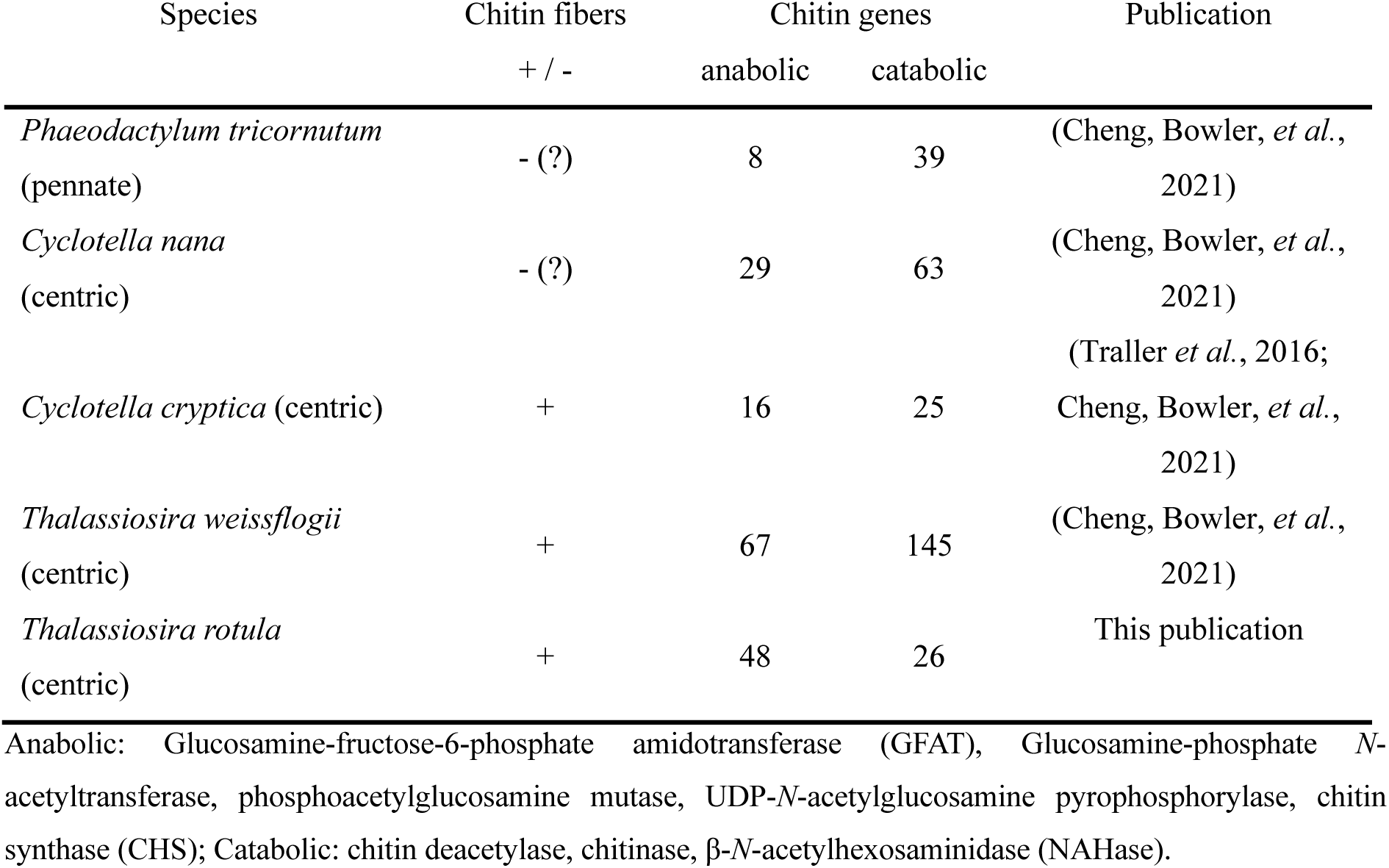
Comparison of chitin metabolism in *Thalassiosirales*. Five different species classified into centric or pennate, and if they extrude chitin fibers.

### Iminosugars shift key metabolic pathways in T.rotula

Iminosugars are carbohydrate analogues where the endocyclic oxygen is replaced by nitrogen, allowing them to act as non-metabolizable mimics of native sugars inhibiting a wide range of carbohydrate active enzymes (CAZy) including chitinases and chitin synthases (Gautier-Lefebvre *et al*., 2005b; Behr, 2011). Light microscopy experiments of *T.rotula* revealed that sub-lethal exposure to synthetic isoleucine-derived iminosugars modulated chitin synthesis and, unexpectedly, produced longer chitin fibers (Holzwarth *et al*., 2022). In contrast, CHS inhibitor Nikkomycin Z reliably shortens fibers over time (unpublished data). Because this contradicts the typical inhibitory nature of iminosugars, we assumed transcriptional reprogramming of the chitin metabolism. An optimized synthesis of the iminosugars gave the products in comparable yields to earlier studies (Holzwarth *et al*., 2022), while reducing the required workup steps and increasing the scale from 0.11 mmol of ImCC to 1.64 mmol, providing sufficient material for transcriptomic profiling of cell cultures. Although iminosugar-treated cells did not show morphological abnormalities, their global metabolic gene expression was significantly reshaped.

ImOH and ImF induced a larger set of differentially expressed genes than ImCC, whose transcriptional profile resembled the DMSO-only control. This likely reflects different recognition and processing of iminosugars by sugar sensing and sugar transporter systems. Plants and algae possess distinct sensing systems for individual sugars, most notably glucose, fructose, and trehalose by hexokinases, sugar transporters and other intrinsic regulatory factors (Rolland, Baena-Gonzalez and Sheen, 2006; Li and Zhao, 2024). Many transporters and kinases display broad substrate specificity (Andersen *et al*., 2025), for example several galactokinases and *N*-acetyl hexosamine kinases can bind (poly-)fluorinated monosaccharides (Keenan *et al*., 2020) and an *Arabidopsis thaliana* transporter even binds the iminosugar 1-deoxynojirimycin (DNJ) (Park *et al*., 2023). Presumably, the different molecular structure of ImCC vs ImOH/ImF leads to alternative uptake or processing mechanisms, which might explain its lower number of DEGs at the same used concentration. Consequently, this study focused on ImOH and ImF treatments.

It was initially anticipated that iminosugars specifically alter expression of chitin-related genes because of the observed morphological effects on the produced chitin fibers (Holzwarth *et al*., 2022). However, unexpectedly, the compounds exerted broad effects on global metabolism. Therefore, genes involved in chitin synthesis, together with key carbohydrate pathways (photosynthesis, Calvin cycle, TCA cycle, glycolysis/gluconeogenesis) and nitrogen metabolism including the diatom-specific OUC were examined. Additionally, ribosome-related genes were investigated due to pronounced upregulation after ImOH and ImF exposure.

The abundance of carbohydrates strongly regulates photosynthesis and central energy stores in algae which can affect their trophic lifestyle (Roth *et al*., 2019; Kurniawan *et al*., 2025). In green alga *Chromochloris zofingiensis*, supplementation of glucose induced transition from photoautotrophy to heterotrophy coinciding with broad downregulation of photosynthetic genes and an increased accumulation of TAGs and starch (Roth *et al*., 2019). Similarly, iminosugar supplementation in *T.rotula* led to downregulation of photosynthetic genes, the diatom-specific FCPs were predominantly repressed.

Chitin biosynthesis is initiated with glutamine transfer onto F6P which is derived from glyceraldehyde-3-phosphate produced in the Calvin cycle or during glycolysis/gluconeogenesis from phosphoenolpyruvate. These pathways are intricately connected with the TCA cycle and the fatty acid metabolism and in diatoms additionally with the OUC (Allen *et al*., 2011; Launay *et al*., 2020). Fig. **5** showed that ImOH (and similarly ImF) reshapes this central metabolism in *T.rotula* by repressing the expression of most of the genes of the Calvin cycle except for RuBisCO (EC 4.1.1.39) and sedoheptulose-bisphosphatase (EC 3.1.3.37) and all genes of glycolysis/gluconeogenesis. Another important metabolite in diatoms is the β-1,3-linked glucan chrysolaminarin that presents the central carbon storage in diatoms (Kwasiborski *et al*., 2025). No DEGs involved in chrysolaminarin biosynthesis could be identified after iminosugar exposure. However, multiple glucanase isoforms were downregulated but due to lack of experimental validation it cannot be verified whether the glucanases are chrysolaminarin-related. In contrast to repressing carbohydrate-related pathways, there seems to be a flux towards fatty acid synthesis after ImOH exposure. Upregulation of genes encoding several enzymes involved in the biosynthesis of (very) long chain fatty acids under iminosugar treatments was observed. Moreover, a gene coding for DGAT, which generates TAGs was found highly upregulated after ImOH exposure. The regulatory networks that decide whether cells allocate their energy towards generation of chrysolaminarin or TAGs is highly dependent on nutrient status (Kwasiborski *et al*., 2025). In *P.tricornutum* TAG accumulates under nutrient supplementation or when chrysolaminarin biosynthesis factors are inhibited (Daboussi *et al*., 2014; Villanova *et al*., 2017).

Furthermore, ImOH (and ImF) affected nitrogen metabolism and the OUC. Downregulation of nitrate-and nitrite-reductase transcripts points to a reduced necessity of extracellular nitrates for assimilation (Fig. **5**). Upregulation of a gene encoding carbamoyl phosphate synthase suggests flux from intracellular ammonia to synthesis of organonitrogen compounds such as amino acids (e.g., arginine and proline biosynthesis or the glutamine and glutamate metabolism) by the OUC. Argininosuccinate lyase gene upregulation suggests feedback to TCA cycle which finally connects nitrogen assimilation to the fatty acid biosynthesis (Albaugh, Mukherjee and Barbul, 2017).

Next to reprogramming of the central carbon and nitrogen metabolism, ImOH and ImF caused upregulation of genes in the pathways ‘ribosome’ and ‘ribosome biogenesis in eukaryotes’ (Fig. **3**). Ribosome biogenesis is one of the most energy costly processes in the cell and represents a central hub for stress response in plants to biotic and abiotic stressors (Moin *et al*., 2016; Garcia-Molina *et al*., 2020; Shore and Albert, 2022; Yang *et al*., 2024). Consistent with previous reports of iminosugars modulating transcription of ribosomal processes, for example β-1-C-propyl-1,4-dideoxy-1,4-imino-L-arabinitol (PDIA) in *Staphylococcus aureus* (Tomusiak-Plebanek *et al*., 2025), DNJ in eri-silkworm (*Samia cynthia ricini*) (Zhang *et al*., 2018), and *N*-(9-methoxynonyl)-DNJ in dengue virus-infected human macrophages (Sayce *et al*., 2021), induction of ribosomal processes including upregulation of genes encoding ribosomal processing proteins, assembly factors and tRNA ligases was observed in the *T.rotula* transcriptome (Fig. **4**, Table **S9**, **S10**). This is indicative of a stress response which leads to increased ribosomal biogenesis.

Overall, this suggests that iminosugars might be imported into *T.rotula* by promiscuous hexokinases or sugar transporters. The iminosugars simulate a pseudo nutrient-rich environment which could trigger a rapid shift from photo-autotrophic to heterotrophic growth by repression of photosynthetic pathways similar to glucose supplementation to *C.zofingiensis* (Roth *et al*., 2019). Concurrently, this coincides with upregulation of parts of the TCA cycle and OUC, thus increasing TAG and organonitrogen synthesis. Potentially as a stress response to the non-metabolizable iminosugars, the cells allocate resources towards ribosome biogenesis (Moin *et al*., 2016; Garcia-Molina *et al*., 2020; Yang *et al*., 2024).

### Chitin biosynthesis under ImOH exposure

Next to reprogramming of central pathways, ImOH exposure affected gene expression related to chitin biosynthesis and its degradation. In *T.rotula*, most of the identified chitin-related genes were predicted to be cytosolic but there was a significant number of ER-targeted genes indicating they are secreted into the extracellular environment or to plasma membranes. Some genes were even predicted to have an SA type-II which embeds proteins into the membrane and prevents cleavage (Gschloessl, Guermeur and Cock, 2008). Generally, expression of genes encoding chitin-degradation factors, especially chitinases, were more repressed while anabolic factors were only slightly repressed.

Three of six GFAT transcripts in *T.rotula* were predicted to be mitochondria-targeted (Fig. **7a**). It is unclear why GFATs might be directed to the mitochondria because it is assumed that glucosamine synthesis is performed in the cytosol. However, GFAT might directly bind mitochondrial glutamine (Bromke, 2013) or F6P might be transported to the mitochondria via transporters. Notably, this phenomenon was observed in *C.cryptica* as well suggesting mitochondrial targeting of GFAT in diatoms might be more common (Traller *et al*., 2016). ImOH was able to significantly repress three GFATs of which two were mitochondria targeted and one ER-targeted. Chitin synthesis is performed by chitin synthases (EC 2.4.1.16). These can be separated into three major divisions (Li *et al*., 2016). Pennate diatom (e.g. *P.tricornutum*) CHS belong predominantly to division 1, while division 2 CHS are characteristic for centric diatoms (Shao *et al*., 2023). The evolutionary history of CHS is complex involving independently formed paralogous groups and multiple horizontal gene transfers explaining the large CHS gene repertoire in some diatoms (Gonçalves *et al*., 2016). In *T.rotula*, 36 out of 42 CHS transcripts were predicted to be cytosolic, five contained SA type-II and two were released to the ER for secretion. ImOH repressed six cytosolic CHS, of which five had a cytochrome b5 (cyt-b5) domain, a conserved tail-anchored membrane protein usually serving as an NADH/NADPH-dependent electron shuttle (Gonçalves *et al*., 2016; Liu, 2022). The lack of heme coordinating histidine residues, however, suggest no redox activity for CHS-fused cyt-b5 and instead it was proposed to bind lipid ligands (Mifsud and Bateman, 2002).

Similar to chitinases in *C.nana* (Cheng, Shao, *et al*., 2021) and *T.weissflogii* (Cheng *et al*., 2025), roughly half (11) of the chitinases were localized to the ER secretory pathway. Additionally, nine were cytosolic and three were predicted to harbor a SA type-II domain indicating some chitinases are anchored to membranes and are not cleaved. The role of chitin degradation in diatoms is not clear but the diversity of the domain architecture suggests adaptation of chitinases to multiple functions which might include remodeling of chitin-associated silica cell walls and extracellular chitin fibers, or as immune defense by recognizing pathogen-associated molecular patterns (Durkin, Mock and Armbrust, 2009; Vaghela *et al*., 2022). In *T.rotula*, we found a total of 10 different domains including the glycosyl hydrolase families GH18 and GH19 (Fig. **7e**, **S6**). Five GH18 chitinases contained an LPMO10 domain. Lytic polysaccharide monooxygenases (LPMOs) are copper enzymes catalyzing oxidative cleavage of recalcitrant chitin. Moreover, they are able to improve chitinase activity by opening the crystalline chitin structure to chitinases (Forsberg *et al*., 2016). These might be involved in degradation of extracellular chitin fibers. The most downregulated chitinases in ImOH were both cytosolic and contained a GH19 domain with TROT011391 possessing both GH18 and GH19 functionality. An ER-targeted NAHase was found to be strongly downregulated as well. NAHases primarily degrade chito oligosaccharides but some show catalytic activity towards chitin (Qu *et al*., 2021). Gene expression of UDP-GlcNAc transporters were also found significantly altered in three instances. Two were slightly downregulated and interestingly, one of the transporters was the only upregulated chitin-related gene under ImOH exposure. Little is known of the impact of GlcNAc transporters on chitin metabolism, however, there is evidence for a role in chitin biosynthesis in some parasites (Nayak and Ghosh, 2019).

With the evidence collected on the transcriptomic level, we find that ImOH exposure correlates with significant repression of genes encoding chitin degrading factors. We therefore assume that the increase of chitin fiber length, as observed previously in *T.rotula* cells (Holzwarth *et al*., 2022), might be in part due to significant repression of genes encoding chitinases and β-*N*-acetyl hexosaminidases. However, this requires additional experimental evidence.

## Acknowledgements

This work was generously financially supported by the Carl Zeiss Foundation for the infrastructure project ChitinFluid (project no. P2019-02004). We thank M. Claußen from the AWI, Germany, who generously provided us with the *T.rotula* cell culture. We also want to thank B. Voss for his invaluable expertise on evaluation of the transcriptomic data sets.

## Author contributions

J.L. and I.M.W. developed the conceptualization, T.W. prepared the iminosugars for the current study, J.L. carried out the experiments with *T.rotula*, investigated the resulting data and performed the formal analysis, J.L. wrote the initial draft of the manuscript and prepared the figures with contributions of T.W., S.L. supervised the synthesis and characterization of iminosugars, I.M.W. supervised and administrated the project, S.L. and I.M.W. acquired the funding. All authors read and approved the final manuscript.

## Data availability

All raw RNA-seq data generated in this work was deposited in the National Center for Biotechnology Information (NCBI) as BioProject PRJNA1482815.

## References

Albaugh, V.L., Mukherjee, K. and Barbul, A. (2017) “Proline Precursors and Collagen Synthesis: Biochemical Challenges of Nutrient Supplementation and Wound Healing,” The Journal of Nutrition, 147(11), pp. 2011–2017. Available at: 10.3945/jn.117.256404.

Allen, A.E. et al. (2011) “Evolution and metabolic significance of the urea cycle in photosynthetic diatoms,” Nature, 473(7346), pp. 203–207. Available at: 10.1038/nature10074.

Andersen, C.G. et al. (2025) “Comparative analysis of STP6 and STP10 unravels molecular selectivity in sugar transport proteins,” Proceedings of the National Academy of Sciences, 122(17), p. e2417370122. Available at: 10.1073/pnas.2417370122.

Barker, T.J. and Boger, D.L. (2012) “Fe(III)/NaBH4-Mediated Free Radical Hydrofluorination of Unactivated Alkenes,” Journal of the American Chemical Society, 134(33), pp. 13588–13591. Available at: 10.1021/ja3063716.

Becker, S., et al. (2017) “Accurate Quantification of Laminarin in Marine Organic Matter with Enzymes from Marine Microbes,” Applied and Environmental Microbiology. Edited by C. Vieille, 83(9), pp. e03389–16. Available at: 10.1128/AEM.03389-16.

Behr, J.-B. (2011) “Chitin Synthase, a Fungal Glycosyltransferase that Is a Valuable Antifungal Target,” CHIMIA, 65(1–2), p. 49. Available at: 10.2533/chimia.2011.49.

Beier, S. and Bertilsson, S. (2013) “Bacterial chitin degradation—mechanisms and ecophysiological strategies,” Frontiers in Microbiology, 4. Available at: 10.3389/fmicb.2013.00149.

Berges, J.A., Franklin, D.J. and Harrison, P.J. (2001) “Evolution of an Artificial Seawater Medium: Improvements in Enriched Seawater, Artificial Water Over the Last Two Decades,” Journal of Phycology, 37(6), pp. 1138–1145. Available at: 10.1046/j.1529-8817.2001.01052.x.

Blackwell, J., Parker, K.D. and Rudall, K.M. (1967) “Chitin fibres of the diatoms Thalassiosira fluviatilis and Cyclotella cryptica,” Journal of Molecular Biology, 28(2), pp. 383–385. Available at: 10/cz7w6d.

Bowler, C. et al. (2008) “The Phaeodactylum genome reveals the evolutionary history of diatom genomes,” Nature, 456(7219), pp. 239–244. Available at: 10.1038/nature07410.

Bromke, M. (2013) “Amino Acid Biosynthesis Pathways in Diatoms,” Metabolites, 3(2), pp. 294–311. Available at: 10.3390/metabo3020294.

Bryłka, K. et al. (2024) “The Cretaceous Diatom Database: A tool for investigating early diatom evolution,” Journal of Phycology, 60(5), pp. 1090–1104. Available at: 10.1111/jpy.13499.

Buchholz, C.R. and Pomerantz, W.C.K. (2021) “19F NMR viewed through two different lenses: ligand-observed and protein-observed 19F NMR applications for fragment-based drug discovery,” RSC Chemical Biology, 2(5), pp. 1312–1330. Available at: 10.1039/d1cb00085c.

Chen, S. et al. (2018) “fastp: an ultra-fast all-in-one FASTQ preprocessor,” Bioinformatics, 34(17), pp. i884–i890. Available at: 10.1093/bioinformatics/bty560.

Cheng, H., Bowler, C., et al. (2021) “Full-Length Transcriptome of Thalassiosira weissflogii as a Reference Resource and Mining of Chitin-Related Genes,” Marine Drugs, 19(7), p. 392. Available at: 10.3390/md19070392.

Cheng, H., Shao, Z., et al. (2021) “Genome-wide identification of chitinase genes in Thalassiosira pseudonana and analysis of their expression under abiotic stresses,” BMC Plant Biology, 21, p. 87. Available at: 10.1186/s12870-021-02849-2.

Cheng, M. et al. (2025) “Genome-Wide Mining of Chitinase Diversity in the Marine Diatom Thalassiosira weissflogii and Functional Characterization of a Novel GH19 Enzyme,” Marine Drugs, 23(4), p. 144. Available at: 10.3390/md23040144.

Conforti, I. and Marra, A. (2021) “Iminosugars as glycosyltransferase inhibitors,” Organic & Biomolecular Chemistry, 19(25), pp. 5439–5475. Available at: 10.1039/D1OB00382H.

Daboussi, F. et al. (2014) “Genome engineering empowers the diatom Phaeodactylum tricornutum for biotechnology,” Nature Communications, 5(1), p. 3831. Available at: 10.1038/ncomms4831.

Di Costanzo, F. et al. (2024) “Thalassiosira rotula,” 2. Available at: 10.17632/c24hb3w4y2.2.

Di Costanzo, F. et al. (2025) “High-quality genome assembly and annotation of Thalassiosira rotula (synonym of Thalassiosira gravida),” Scientific Data, 12(1), p. 310. Available at: 10.1038/s41597-025-04634-4.

Downey, K.M. et al. (2023) “The dynamic response to hypo-osmotic stress reveals distinct stages of freshwater acclimation by a euryhaline diatom,” Molecular Ecology, 32(11), pp. 2766–2783. Available at: 10.1111/mec.16703.

Durkin, C.A., Mock, T. and Armbrust, E.V. (2009) “Chitin in Diatoms and Its Association with the Cell Wall,” Eukaryotic Cell, 8(7), pp. 1038–1050. Available at: 10.1128/EC.00079-09.

Dyhrman, S.T. et al. (2012) “The Transcriptome and Proteome of the Diatom Thalassiosira pseudonana Reveal a Diverse Phosphorus Stress Response,” PLOS ONE, 7(3), p. e33768. Available at: 10.1371/journal.pone.0033768.

Dyrløv Bendtsen, J., et al. (2004) “Improved Prediction of Signal Peptides: SignalP 3.0,” Journal of Molecular Biology, 340(4), pp. 783–795. Available at: 10.1016/j.jmb.2004.05.028.

Ehrlich, H. and Witkowski, A. (2015) “Biomineralization in Diatoms: The Organic Templates,” pp. 39–58. Available at: 10.1007/978-94-017-9398-8_3.

Field, C.B. et al. (1998) “Primary Production of the Biosphere: Integrating Terrestrial and Oceanic Components,” Science, 281(5374), pp. 237–240. Available at: 10.1126/science.281.5374.237.

Forsberg, Z. et al. (2016) “Structural and Functional Analysis of a Lytic Polysaccharide Monooxygenase Important for Efficient Utilization of Chitin in *Cellvibrio japonicus*\*,” Journal of Biological Chemistry, 291(14), pp. 7300–7312. Available at: 10.1074/jbc.M115.700161.

Garcia-Molina, A. et al. (2020) “Translational Components Contribute to Acclimation Responses to High Light, Heat, and Cold in *Arabidopsis*,” iScience, 23(7), p. 101331. Available at: 10.1016/j.isci.2020.101331.

Gautier-Lefebvre, I. et al. (2005a) “Iminosugars as glycosyltransferase inhibitors: synthesis of polyhydroxypyrrolidines and their evaluation on chitin synthase activity,” European Journal of Medicinal Chemistry, 40(12), pp. 1255–1261. Available at: 10/ftsfzf.

Gautier-Lefebvre, I. et al. (2005b) “Iminosugars as glycosyltransferase inhibitors: synthesis of polyhydroxypyrrolidines and their evaluation on chitin synthase activity,” European Journal of Medicinal Chemistry, 40(12), pp. 1255–1261. Available at: 10.1016/j.ejmech.2005.07.001.

Gimenez, D. et al. (2021) “19F NMR as a tool in chemical biology,” Beilstein Journal of Organic Chemistry, 17(1), pp. 293–318. Available at: 10.3762/bjoc.17.28.

Gonçalves, I.R. et al. (2016) “Genome-wide analyses of chitin synthases identify horizontal gene transfers towards bacteria and allow a robust and unifying classification into fungi,” BMC Evolutionary Biology, 16(1), p. 252. Available at: 10.1186/s12862-016-0815-9.

Gruber, A. et al. (2025) “ASAFind 2.0: multi-class protein targeting prediction for diatoms and algae with complex plastids,” The Plant Journal: For Cell and Molecular Biology, 122(5), p. e70138. Available at: 10.1111/tpj.70138.

Gschloessl, B., Guermeur, Y. and Cock, J.M. (2008) “HECTAR: A method to predict subcellular targeting in heterokonts,” BMC Bioinformatics, 9(1), p. 393. Available at: 10.1186/1471-2105-9-393.

Gügi, B. et al. (2015) “Diatom-Specific Oligosaccharide and Polysaccharide Structures Help to Unravel Biosynthetic Capabilities in Diatoms,” Marine Drugs, 13(9), pp. 5993–6018. Available at: 10/gk6znn.

Harrison, P.J., Waters, R.E. and Taylor, F.J.R. (1980) “A Broad Spectrum Artificial Sea Water Medium for Coastal and Open Ocean Phytoplankton1,” Journal of Phycology, 16(1), pp. 28–35. Available at: 10.1111/j.0022-3646.1980.00028.x.

Herth, W. (1979) “The site of β-chitin fibril formation in centric diatoms. II. The chitin-forming cytoplasmic structures,” Journal of Ultrastructure Research, 68(1), pp. 16–27. Available at: 10.1016/s0022-5320(79)90138-2.

Holzwarth, M. et al. (2022) “Modulating chitin synthesis in marine algae with iminosugars obtained by SmI2 and FeCl3-mediated diastereoselective carbonyl ene reaction,” Organic & Biomolecular Chemistry, 20, pp. 6606–6618. Available at: 10.1039/D2OB00907B.

Jones, J.C. et al. (2021) “Soluble Methane Monooxygenase Component Interactions Monitored by 19F NMR,” Biochemistry, 60(25), pp. 1995–2010. Available at: 10.1021/acs.biochem.1c00293.

Jones, P. et al. (2014) “InterProScan 5: genome-scale protein function classification,” Bioinformatics, 30(9), pp. 1236–1240. Available at: 10.1093/bioinformatics/btu031.

Keenan, T. et al. (2020) “Profiling Substrate Promiscuity of Wild-Type Sugar Kinases for Multi-fluorinated Monosaccharides,” Cell Chemical Biology, 27(9), pp. 1199–1206.e5. Available at: 10.1016/j.chembiol.2020.06.005.

Kim, D. et al. (2019) “Graph-based genome alignment and genotyping with HISAT2 and HISAT-genotype,” Nature Biotechnology, 37(8), pp. 907–915. Available at: 10.1038/s41587-019-0201-4.

Kumar, K. et al. (2025) “Simultaneous assessment of membrane bilayer structure and drug insertion by 19F solid-state NMR,” Biophysical Journal, 124(2), pp. 256–266. Available at: 10.1016/j.bpj.2024.11.3319.

Kurniawan, S.B. et al. (2025) “Autotrophic vs. heterotrophic microalgae: Juxtaposition of performances in treating organic-rich effluent,” Desalination and Water Treatment, 322, p. 101159. Available at: 10.1016/j.dwt.2025.101159.

Kwasiborski, A. et al. (2025) “Chrysolaminarin metabolism in diatoms: Pathways, regulation, and biotechnological perspectives,” Journal of Applied Phycology, 37(6), pp. 3993–4006. Available at: 10.1007/s10811-025-03695-7.

Launay, H. et al. (2020) “Regulation of Carbon Metabolism by Environmental Conditions: A Perspective From Diatoms and Other Chromalveolates,” Frontiers in Plant Science, 11. Available at: 10.3389/fpls.2020.01033.

Lebeau, T. and Robert, J.-M. (2003) “Diatom cultivation and biotechnologically relevant products. Part II: current and putative products,” Applied Microbiology and Biotechnology, 60(6), pp. 624–632. Available at: 10.1007/s00253-002-1177-3.

Li, G. and Zhao, Y. (2024) “The critical roles of three sugar-related proteins (HXK, SnRK1, TOR) in regulating plant growth and stress responses,” Horticulture Research, 11(6), p. uhae099. Available at: 10.1093/hr/uhae099.

Li, L. et al. (2021) “The Draft Genome of the Centric Diatom Conticribra weissflogii (Coscinodiscophyceae, Ochrophyta),” Protist, p. 125845. Available at: 10/gnx9n3.

Li, M. et al. (2016) “Evolution and Functional Insights of Different Ancestral Orthologous Clades of Chitin Synthase Genes in the Fungal Tree of Life,” Frontiers in Plant Science, 7. Available at: 10.3389/fpls.2016.00037.

Liao, Y., Smyth, G.K. and Shi, W. (2014) “featureCounts: an efficient general purpose program for assigning sequence reads to genomic features,” Bioinformatics, 30(7), pp. 923–930. Available at: 10.1093/bioinformatics/btt656.

Liu, C.-J. (2022) “Cytochrome b5: A versatile electron carrier and regulator for plant metabolism,” Frontiers in Plant Science, 13, p. 984174. Available at: 10.3389/fpls.2022.984174.

Love, M.I., Huber, W. and Anders, S. (2014) “Moderated estimation of fold change and dispersion for RNA-seq data with DESeq2,” Genome Biology, 15(12), p. 550. Available at: 10.1186/s13059-014-0550-8.

Ludwig, J. et al. (2025) “Ro(a)d to New Functional Materials: Sustainable Isolation of High-Aspect-Ratio β-Chitin Microrods from Marine Algae,” Bioengineering, 12(9), p. 969. Available at: 10.3390/bioengineering12090969.

Marella, T.K., Bhattacharjya, R. and Tiwari, A. (2021) “Impact of organic carbon acquisition on growth and functional biomolecule production in diatoms,” Microbial Cell Factories, 20(1), p. 135. Available at: 10.1186/s12934-021-01627-x.

Martin-Jézéquel, V., Hildebrand, M. and Brzezinski, M.A. (2000) “Silicon Metabolism in Diatoms: Implications for Growth,” Journal of Phycology, 36(5), pp. 821–840. Available at: 10.1046/j.1529-8817.2000.00019.x.

McLachlan, J., McInnes, A.G. and Falk, M. (1965) “Studies on the chitan (chitin: poly-n-acetylglucosamine) fibers of the diatom thalassiosira fluviatilis hustedt: i production and isolation of chitan fibers,” Canadian Journal of Botany, 43(6), pp. 707–713. Available at: 10.1139/b65-079.

Mifsud, W. and Bateman, A. (2002) “Membrane-bound progesterone receptors contain a cytochrome b5-like ligand-binding domain,” Genome Biology, 3(12), p. research0068.1. Available at: 10.1186/gb-2002-3-12-research0068.

Moin, M. et al. (2016) “Rice Ribosomal Protein Large Subunit Genes and Their Spatio-temporal and Stress Regulation,” Frontiers in Plant Science, 7. Available at: 10.3389/fpls.2016.01284.

Murison, V. et al. (2025) “Diatom triacylglycerol metabolism: from carbon fixation to lipid droplet degradation,” Biological Reviews, 100(4), pp. 1423–1443. Available at: 10.1111/brv.70006.

Nayak, S. and Ghosh, S.K. (2019) “Nucleotide sugar transporters of Entamoeba histolytica and Entamoeba invadens involved in chitin synthesis,” Molecular and Biochemical Parasitology, 234, p. 111224. Available at: 10.1016/j.molbiopara.2019.111224.

Park, J. et al. (2023) “Exploring the Substrate Specificity of a Sugar Transporter with Biosensors and Cheminformatics,” ACS Synthetic Biology, 12(2), pp. 565–571. Available at: 10.1021/acssynbio.2c00571.

Paysan-Lafosse, T. et al. (2025) “The Pfam protein families database: embracing AI/ML,” Nucleic Acids Research, 53(D1), pp. D523–D534. Available at: 10.1093/nar/gkae997.

Pertea, M. et al. (2015) “StringTie enables improved reconstruction of a transcriptome from RNA-seq reads,” Nature Biotechnology, 33(3), pp. 290–295. Available at: 10.1038/nbt.3122.

Prihoda, J. et al. (2012) “Chloroplast-mitochondria cross-talk in diatoms,” Journal of Experimental Botany, 63(4), pp. 1543–1557. Available at: 10.1093/jxb/err441.

Qu, T., et al. (2021) “A Novel GH Family 20 β-N-acetylhexosaminidase With Both Chitosanase and Chitinase Activity From Aspergillus oryzae,” Frontiers in Molecular Biosciences, 8. Available at: 10.3389/fmolb.2021.684086.

Rogg, L.E. et al. (2012) “Regulation of expression, activity and localization of fungal chitin synthases,” Medical mycology: official publication of the International Society for Human and Animal Mycology, 50(1), pp. 2–17. Available at: 10.3109/13693786.2011.577104.

Rolland, F., Baena-Gonzalez, E. and Sheen, J. (2006) “Sugar sensing and signaling in plants: conserved and novel mechanisms,” Annual Review of Plant Biology, 57, pp. 675–709. Available at: 10.1146/annurev.arplant.57.032905.105441.

Roth, M.S. et al. (2019) “Regulation of Oxygenic Photosynthesis during Trophic Transitions in the Green Alga *Chromochloris zofingiensis*,” The Plant Cell, 31(3), pp. 579–601. Available at: 10.1105/tpc.18.00742.

Safadi, R. et al. (2025) “Structural organization of the organic sheath that delineates extracellular seta silicification in diatoms,” Journal of Structural Biology, 217(2), p. 108205. Available at: 10.1016/j.jsb.2025.108205.

Sayce, A.C. et al. (2021) “Pathogen-induced inflammation is attenuated by the iminosugar M *O* N-DNJ via modulation of the unfolded protein response,” Immunology, 164(3), pp. 587–601. Available at: 10.1111/imm.13393.

Schönitzer, V. and Weiss, I.M. (2007) “The structure of mollusc larval shells formed in the presence of the chitin synthase inhibitor Nikkomycin Z,” BMC structural biology, 7, p. 71. Available at: 10.1186/1472-6807-7-71.

Shao, Z. et al. (2019) “Comparative characterization of putative chitin deacetylases from *Phaeodactylum tricornutum* and *Thalassiosira pseudonana* highlights the potential for distinct chitin-based metabolic processes in diatoms,” New Phytologist, 221(4), pp. 1890–1905. Available at: 10.1111/nph.15510.

Shao, Z. et al. (2023) “Characterization of a Marine Diatom Chitin Synthase Using a Combination of Meta-Omics, Genomics, and Heterologous Expression Approaches,” mSystems, 0(0), pp. e01131–22. Available at: 10.1128/msystems.01131-22.

Shore, D. and Albert, B. (2022) “Ribosome biogenesis and the cellular energy economy,” Current Biology, 32(12), pp. R611–R617. Available at: 10.1016/j.cub.2022.04.083.

Sievers, F. et al. (2011) “Fast, scalable generation of high-quality protein multiple sequence alignments using Clustal Omega,” Molecular Systems Biology, 7(1), p. MSB201175. Available at: 10.1038/msb.2011.75.

Souza, C.P. et al. (2011) “The Importance of Chitin in the Marine Environment,” Marine Biotechnology, 13(5), pp. 823–830. Available at: 10.1007/s10126-011-9388-1.

Stiller, J.W. et al. (2014) “The evolution of photosynthesis in chromist algae through serial endosymbioses,” Nature Communications, 5(1), p. 5764. Available at: 10.1038/ncomms6764.

Supek, F. et al. (2011) “REVIGO Summarizes and Visualizes Long Lists of Gene Ontology Terms,” PLOS ONE, 6(7), p. e21800. Available at: 10.1371/journal.pone.0021800.

Tomusiak-Plebanek, A. et al. (2025) “Transcriptomic Analysis of Biofilm Formation Inhibition by PDIA Iminosugar in Staphylococcus aureus,” Antibiotics, 14(7), p. 668. Available at: 10.3390/antibiotics14070668.

Traller, J.C. et al. (2016) “Genome and methylome of the oleaginous diatom Cyclotella cryptica reveal genetic flexibility toward a high lipid phenotype,” Biotechnology for Biofuels, 9(1), p. 258. Available at: 10.1186/s13068-016-0670-3.

Tréguer, P. et al. (2018) “Influence of diatom diversity on the ocean biological carbon pump,” Nature Geoscience, 11(1), pp. 27–37. Available at: 10.1038/s41561-017-0028-x.

Vaghela, B. et al. (2022) “Plant chitinases and their role in plant defense: A comprehensive review,” Enzyme and Microbial Technology, 159, p. 110055. Available at: 10.1016/j.enzmictec.2022.110055.

Villanova, V. et al. (2017) “Investigating mixotrophic metabolism in the model diatom *Phaeodactylum tricornutum*,” Philosophical Transactions of the Royal Society B: Biological Sciences, 372(1728), p. 20160404. Available at: 10.1098/rstb.2016.0404.

Villar, E. et al. (2018) “The Ocean Gene Atlas: exploring the biogeography of plankton genes online,” Nucleic Acids Research, 46(W1), pp. W289–W295. Available at: 10.1093/nar/gky376.

Wagner, H. et al. (2017) “Towards an understanding of the molecular regulation of carbon allocation in diatoms: the interaction of energy and carbon allocation,” Philosophical Transactions of the Royal Society B: Biological Sciences, 372(1728), p. 20160410. Available at: 10.1098/rstb.2016.0410.

Waterhouse, A.M., et al. (2009) “Jalview Version 2—a multiple sequence alignment editor and analysis workbench,” Bioinformatics, 25(9), pp. 1189–1191. Available at: 10.1093/bioinformatics/btp033.

Wegmann, M. et al. (2026) “Structural and chemical properties of insect’s chitin-containing extracellular matrices,” Insect Biochemistry and Molecular Biology, 193, p. 104608. Available at: 10.1016/j.ibmb.2026.104608.

Wickham, H. (2016) ggplot2. Cham: Springer International Publishing (Use R!). Available at: 10.1007/978-3-319-24277-4.

Wilhelm, C. et al. (2006) “The Regulation of Carbon and Nutrient Assimilation in Diatoms is Significantly Different from Green Algae,” Protist, 157(2), pp. 91–124. Available at: 10.1016/j.protis.2006.02.003.

Wustmann, M. et al. (2020) “Chitin synthase localization in the diatom Thalassiosira pseudonana,” BMC Materials, 2(1), p. 10. Available at: 10.1186/s42833-020-00016-9.

Yang, F. et al. (2024) “Benzophenone-4 inhibition in marine diatoms: Physiological and molecular perspectives,” Ecotoxicology and Environmental Safety, 284, p. 117021. Available at: 10.1016/j.ecoenv.2024.117021.

Yu, G. et al. (2012) “clusterProfiler: an R Package for Comparing Biological Themes Among Gene Clusters,” OMICS : a Journal of Integrative Biology, 16(5), pp. 284–287. Available at: 10.1089/omi.2011.0118.

Zhang, S.-Z. et al. (2018) “Comparative transcriptome analysis reveals significant metabolic alterations in eri-silkworm (Samia cynthia ricini) haemolymph in response to 1-deoxynojirimycin,” PLOS ONE, 13(1), p. e0191080. Available at: 10.1371/journal.pone.0191080.

